# Projection-dependent VGluT1 and VGluT2 expression and terminal morphology in mouse visual circuits

**DOI:** 10.64898/2026.08.31.748331

**Authors:** Guillaume Laliberté, Robert Tremblay-Laliberté, Denis Boire

**Author notes:** **Corresponding author: Denis Boire PhD**, Département d’anatomie, Université du Québec à Trois-Rivières 3351, boulevard des Forges, C.P. 500 Trois-Rivières, Québec, G9A 5H7. GL and RTL share first authorship.

## Abstract

Vesicular glutamate transporters 1 and 2 (VGluT1 and VGluT2) exhibit largely complementary distributions and have been proposed as molecular markers of descending modulatory and ascending driver-like pathways, respectively. However, this correspondence has rarely been tested directly in anatomically identified projections. We combined anterograde Phaseolus vulgaris leucoagglutinin tracing with simultaneous VGluT1 and VGluT2 immunofluorescence to characterize glutamatergic boutons arising from the primary and secondary visual cortices, lateral geniculate nucleus, lateral posterior thalamic nucleus, and superior colliculus in adult mice. Among 2,178 singly labelled boutons, VGluT1 predominated in corticothalamic, corticopontine, corticotectal, corticostriatal, and corticocortical feedback projections. Conversely, thalamocortical, tectothalamic, thalamostriatal, and tectopontine projections were almost exclusively VGluT2-positive. Projection direction did not fully predict transporter phenotype: ascending corticocortical projections remained predominantly VGluT1-positive, whereas the descending tectopontine projection was exclusively VGluT2-positive. Only 12 of 2,190 VGluT-immunoreactive boutons exhibited detectable VGluT1/VGluT2 colocalization. Morphometric analyses further revealed an interaction between VGluT isoform and projection direction. Within ascending projections, VGluT2-positive boutons and puncta were larger than their VGluT1-positive counterparts, whereas no isoform-related difference in bouton area was detected within descending projections. These findings demonstrate an association between VGluT phenotype and projection class in the mouse visual system. VGluT isoforms therefore provide informative markers of pathway organization, but neither transporter identity nor terminal size alone constitutes an invariant molecular indicator of projection direction or driver–modulator function.

**Graphical Abstract:** 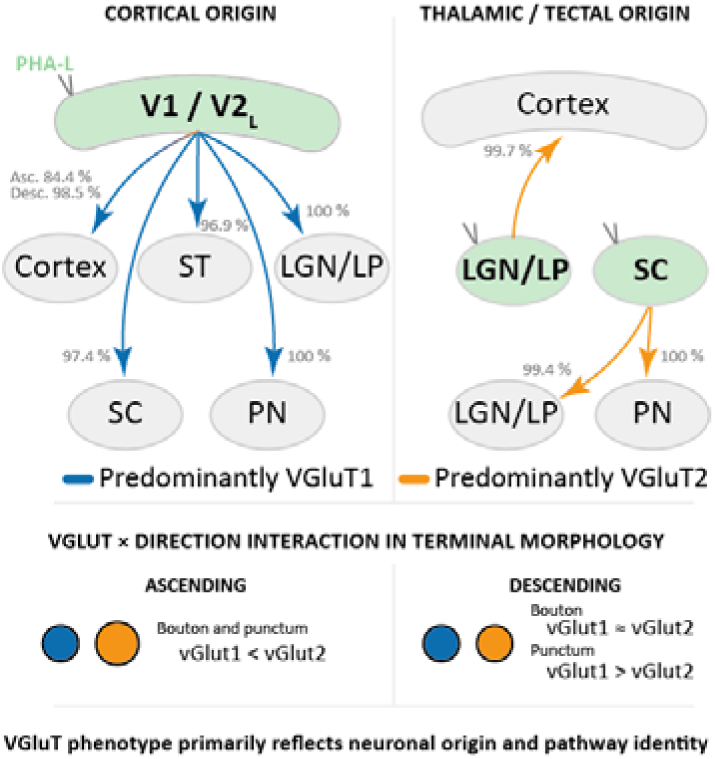

Cortical (V1/V2L) projections are predominantly VGluT1+, whereas thalamic and tectal (LGN/LP/SC) projections are predominantly VGluT2+, linking transporter phenotype to neuronal origin and pathway identity. Terminal morphology shows a VGluT×direction interaction: ascending boutons and puncta are larger for VGluT2, whereas descending pathways show comparable bouton sizes but larger VGluT1+ puncta.

## Introduction

Glutamate is the principal excitatory neurotransmitter in the mammalian central nervous system. Its accumulation into synaptic vesicles is mediated by three vesicular transporters (VGluT1-3), among which VGluT1 and VGluT2 are the predominant isoforms in synapses (Bellocchio et al. 2000; Takamori et al. 2000; Fremeau et al. 2001), whereas VGluT3 was subsequently identified in more restricted neuronal populations (Fremeau et al. 2002). These two predominant isoforms exhibit largely complementary distributions. VGluT1 is predominantly expressed by with cortical, hippocampal, and cerebellar neurons, whereas VGluT2 is in neurons of the thalamus, brainstem, and other subcortical structures (Fremeau *et al*. 2001; Herzog et al. 2001; Kaneko and Fujiyama 2002; Varoqui et al. 2002; Fremeau, Kam, et al. 2004). Nevertheless, this segregation is not absolute and remains incompletely characterized. The coexpression of VGluT1 and VGluT2 mRNAs has been reported in subsets of adult thalamic neurons (Barroso-Chinea et al. 2008). At the protein level, VGluT1 and VGluT2 have been colocalized within individual synapses and synaptic vesicles in the developing mouse hippocampus (Herzog et al. 2006), as well as in a subset of neocortical terminals, with colocalization in the visual cortex being more frequent during postnatal development than in adulthood (Nakamura et al. 2007).

Beyond their anatomical distribution, VGluT1- and VGluT2-expressing terminals differ in several presynaptic properties. VGluT1-expressing synapses are generally associated with a lower initial release probability and less short-term depression, or, in some pathways, greater facilitation, whereas VGluT2-expressing synapses are more frequently associated with a higher release probability and short-term depression (Fremeau, Kam*, et al.* 2004; Fremeau, Voglmaier, et al. 2004; Wojcik et al. 2004; Weston et al. 2011). These properties resemble, respectively, the modulatory and driving modes of glutamatergic transmission described in corticothalamic and thalamocortical circuits. Driver inputs typically produce large, ionotropic receptor-mediated responses and transmit the principal information represented by their target neurons, whereas modulatory inputs regulate the efficacy and temporal properties of this transmission through both ionotropic and metabotropic mechanisms (Sherman and Guillery 1998; Sherman and Guillery 2001; Reichova and Sherman 2004; Sherman and Guillery 2011).

Within sensory systems, this functional distinction has been related to the molecular identity and morphology of glutamatergic terminals. Ascending retinogeniculate and thalamocortical projections commonly express VGluT2 and form comparatively large terminals associated with driver-like transmission (Fujiyama et al. 2003; Reichova and Sherman 2004; Nahmani and Erisir 2005). Conversely, corticothalamic feedback projections predominantly express VGluT1 and typically form smaller terminals exhibiting low initial efficacy and short-term facilitation, consistent with modulatory transmission (Reichova and Sherman 2004; Yoshida et al. 2009; Lindström et al. 2020). Consistent with this organization, VGluT2-immunoreactive terminals associated with driver pathways are often larger than VGluT1-immunoreactive or corticothalamic terminals in specific sensory circuits, particularly in retinogeniculate and geniculocortical pathways (Fujiyama *et al*. 2003; Nahmani and Erisir 2005). Large terminal profiles have also been recognized as a characteristic of several driver pathways independently of VGluT phenotype (Sherman and Guillery 2011; Rovo et al. 2012). Comparative anatomical studies have therefore suggested that VGluT2 is preferentially associated with ascending or feedforward pathways, whereas VGluT1 predominates in descending or feedback pathways of the visual system (Balaram, Hackett, et al. 2011; Balaram et al. 2013; Balaram et al. 2015; Abbas Farishta et al. 2022). These associations broadly parallel the driver–modulator distinction, but transporter identity alone does not establish the functional class of a projection.

Most previous investigations of visual pathways have inferred the projection-specific expression of VGluT isoforms from their regional or laminar distributions or from the localization of VGluT-expressing cell bodies (Balaram, Hackett*, et al.* 2011; Balaram, Takahata, et al. 2011; Rovo *et al*. 2012; Balaram *et al*. 2013; Balaram *et al*. 2015). Direct colocalization of VGluT2 with anatomically identified terminals has been demonstrated in retinogeniculate and geniculocortical projections (Fujiyama *et al*. 2003; Nahmani and Erisir 2005). VGluT2 expression and comparatively large bouton profiles have also been reported for pulvinar projections to visual cortex, consistent with a driver-like organization (Marion et al. 2013). By contrast, the VGluT phenotype of identified corticocortical, corticofugal, tectothalamic, and higher order thalamocortical projections remains incompletely characterized in the mouse. It remains unclear whether VGluT1 and VGluT2 reliably distinguish feedforward from feedback corticocortical projections and whether the two isoforms can coexist within individual terminals of anatomically identified visual pathways.

In the present study, we combined anterograde tracing with simultaneous immunofluorescent detection of VGluT1 and VGluT2 to determine the transporter phenotype of axonal boutons arising from the primary visual cortex (V1), lateral secondary visual cortex (V2_L_), dorsal lateral geniculate nucleus (dLGN), lateral posterior thalamic nucleus (LP), and superior colliculus (SC) in the mouse. We examined corticocortical, corticothalamic, corticotectal, corticopontine, corticostriatal, thalamocortical, thalamostriatal, tectothalamic, and tectopontine projections. We tested whether VGluT2 predominates in ascending projections and VGluT1 in descending projections, and whether VGluT2-positive (VGluT2+) boutons exhibit larger profiles than VGluT1-positive (VGluT1+) boutons, as predicted from previously reported associations between transporter phenotype, projection direction, and terminal morphology. We additionally assessed whether VGluT1 and VGluT2 are coexpressed within individual boutons of identified visual pathways.

## Materials and Methods

### Animals, housing, and surgery

Twenty-six eight-week-old female C57Bl/6J mice weighing 20–23 g were included in the analyses (Charles River Laboratories, Montréal, QC, Canada). Mice were group-housed under institutionally controlled environmental conditions, with target ranges of 20–26°C and 40–60% relative humidity, 15–20 air changes per hour, and a 12h light period. Autoclaved water and irradiated food were available *ad libitum*. Environmental enrichment included nesting material, a shelter, and a gnawing item, in accordance with institutional animal-care procedures. Mice were assigned to one of five tracer-injection sites: the primary visual cortex (V1; n = 7), lateral secondary visual cortex (V2L; n = 5), dorsal lateral geniculate nucleus (dLGN; n = 5), lateral posterior thalamic nucleus (LP; n = 5), or superior colliculus (SC; n = 4).

All procedures were conducted in accordance with the guidelines of the Canadian Council on Animal Care and were approved by the *Comité de bons soins aux animaux of the Université du Québec à Trois-Rivières*. Before surgery, mice received subcutaneous buprenorphine (0.10 mg/kg) and sterile saline (0.5 mL). Anaesthesia was induced with 5% isoflurane in oxygen and maintained with 2% isoflurane throughout the procedure. Ophthalmic lubricant was applied to prevent corneal dehydration, and vital signs and anaesthetic depth were monitored throughout surgery.

Mice were positioned in a stereotaxic apparatus (model 51730U; Stoelting, Wood Dale, IL, USA), and a midline scalp incision was made to expose the skull. A small craniotomy approximately 0.9 mm in diameter was performed above the selected injection site. Stereotaxic coordinates and injection duration for each target are provided in **Table 1**. Unconjugated Phaseolus vulgaris leucoagglutinin (PHA-L; 2.5%; catalog no. L-1110, Vector Laboratories, Newark, CA, USA) was prepared in 0.1 M phosphate-buffered saline and loaded into a glass micropipette with an internal tip diameter of approximately 15 µm. PHA-L was delivered iontophoretically using positive-current pulses of 2 µA with a 7-s on/7-s off duty cycle for 10–20 min (Midgard Precision Current Source; Stoelting).

**TABLE 1.** Injection sites and stereotaxic coordinates (AP anteroposterior, ML, mediolateral and DV dorsoventral depth). Cortical injections were at four depths to distribute tracers to all cortical layers. Injection time was subdivided equally to each depth.

| CASE | AP<br>(MM) | ML<br>(MM) | DV<br>(MM) | ANGLE° | INJECTION<br>TIME<br>(MIN) |
| --- | --- | --- | --- | --- | --- |
| **V1 INJECTION** |  |  |  |  |  |
| **Q8** | -3.50 | 2.25 | 0.25/0.40/0.55/0.70 | 0 | 12 |
| **Q9** | -3.50 | 2.25 | 0.25/0.40/0.55/0.70 | 0 | 12 |
| **Q10** | -3.50 | 2.25 | 0.25/0.40/0.55/0.70 | 0 | 12 |
| **Q11** | -3.50 | 2.25 | 0.25/0.40/0.55/0.70 | 0 | 12 |
| **U1** | -3.50 | 2.25 | 0.25/0.40/0.55/0.70 | 0 | 20 |
| **Y12** | -3.50 | 2.25 | 0.25/0.40/0.55/0.70 | 0 | 20 |
| **Y13** | -3.50 | 2.25 | 0.25/0.40/0.55/0.70 | 0 | 20 |
| ***V2<sub>L</sub> INJECTION*** |  |  |  |  |  |
| **S2** | -3.78 | 3.70 | 0.25/0.40/0.55/0.70 | 30 | 12 |
| **U2** | -3.78 | 3.70 | 0.25/0.40/0.55/0.70 | 30 | 20 |
| **U3** | -3.78 | 3.70 | 0.25/0.40/0.55/0.70 | 30 | 20 |
| **X1** | -3.78 | 3.70 | 0.25/0.40/0.55/0.70 | 30 | 20 |
| **AA1** | -3.78 | 3.70 | 0.25/0.40/0.55/0.70 | 30 | 20 |
| ***DLGN INJECTION*** |  |  |  |  |  |
| **V4** | -2.30 | 2.00 | 2.44 | 0 | 10 |
| **V5** | -2.30 | 2.00 | 2.44 | 0 | 10 |
| **W1** | -2.30 | 2.00 | 2.50 | 0 | 20 |
| **W2** | -2.30 | 2.00 | 2.50 | 0 | 20 |
| **W3** | -2.30 | 2.00 | 2.50 | 0 | 20 |
| ***LP INJECTION*** |  |  |  |  |  |
| **U4** | -1.94 | 1.50 | 2.50 | 0 | 10 |
| **W7** | -1.94 | 1.50 | 2.50 | 0 | 20 |
| **Y9** | -1.94 | 1.50 | 2.50 | 0 | 20 |
| **Y11** | -1.94 | 1.50 | 2.50 | 0 | 20 |
| **Y14** | -1.94 | 1.50 | 2.50 | 0 | 20 |
| ***SC INJECTION*** |  |  |  |  |  |
| **Y6** | -3.50 | 0.50 | 1.40/1.50 | 0 | 20 |
| **Y7** | -3.50 | 0.50 | 1.40/1.50 | 0 | 20 |
| **Y8** | -3.50 | 0.50 | 1.40/1.50 | 0 | 20 |
| **Z4** | -3.50 | 0.50 | 1.40/1.50 | 0 | 20 |

Following iontophoresis, the micropipette was left in place for 5 min to minimize tracer reflux along the injection tract and was then slowly withdrawn. The scalp incision was closed using 6-0 absorbable sutures (Vicryl). Carprofen (10 mg/kg, subcutaneous; Rimadyl) was administered immediately after surgery and once daily for the following two days. Animals were monitored during postoperative recovery according to institutional animal-care procedures.

Injection sites were subsequently verified histologically. Animals were included only when the injection core was located within the intended structure and produced clearly identifiable anterograde labelling in expected projection targets. Animals in which the tracer deposit extended substantially beyond the boundaries of the intended injection site were excluded. Twenty-six animals met these inclusion criteria and were retained for analysis (**Figure 1)**.

**Figure 1.**
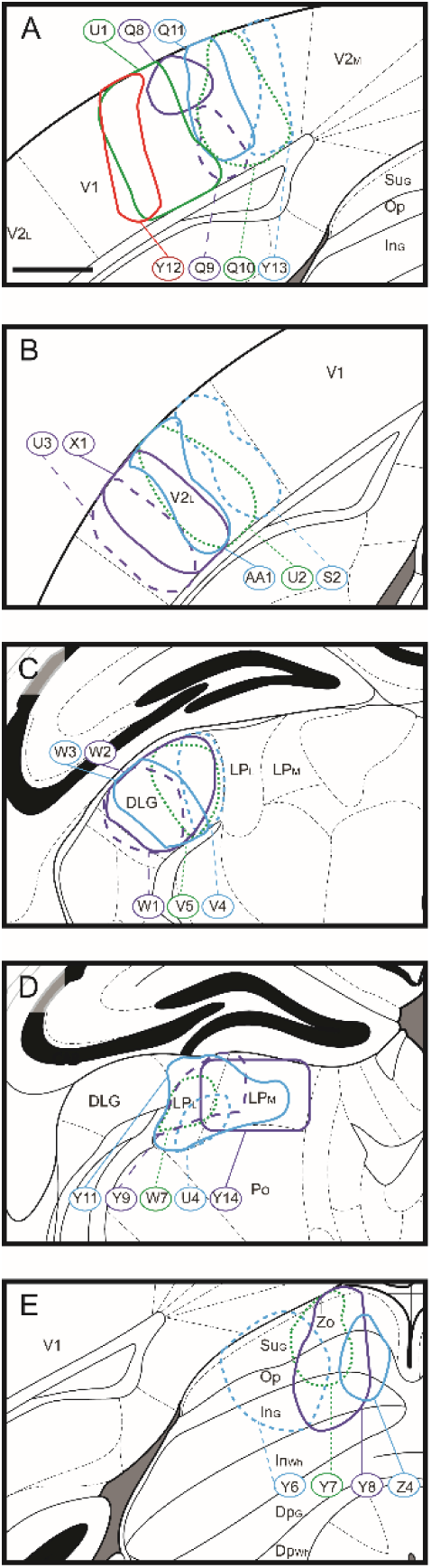
Location and extent of PHA-L injection sites. Reconstructions show the distribution of individual injection sites in the primary visual cortex (V1; **A**), lateral secondary visual cortex (V2_L_; **B**), dorsal lateral geniculate nucleus (DLG; **C**), lateral posterior thalamic nucleus (LP; **D**), and superior colliculus (SC; **E**). Coloured contours represent the reconstructed extent of PHA-L labelling for individual cases; case identifiers correspond to those reported in Table 1. Line colours and patterns are used solely to distinguish overlapping cases. Anatomical boundaries were delineated according to Paxinos and Franklin (2007). (**A-E**) Scale bar = 500 µm.

### Perfusion and tissue preparation

Seven days after PHA-L injection, mice were deeply anaesthetized with 5% isoflurane and transcardially perfused with 0.1 M phosphate-buffered saline (PBS; pH 7.3), followed by 4% paraformaldehyde in 0.1 M PBS. Brains were removed and postfixed in 4% paraformaldehyde for 1 h. They were then cryoprotected overnight at 4°C in 30% sucrose in 0.1 M PBS, until they sank. Cryoprotected brains were stored at −80°C until histologic processing.

Brains were coronally sectioned at a nominal thickness of 40 µm using a freezing microtome (SM2400; Leica Microsystems). Sections were collected sequentially into six series, resulting in a 240 µm interval between consecutive sections within each series. One series from each brain was routinely processed for immunofluorescence. The remaining series were stored frozen for subsequent use.

### Immunofluorescence

Free-floating sections were incubated in 50% methanol for 20 min and washed three times in 0.1 M phosphate-buffered saline (PBS; pH 7.3). Sections were then incubated overnight in a blocking and permeabilization solution containing 2% normal donkey serum and 2% Triton X- 100 in 0.1 M PBS. Subsequently, sections were incubated for 72 h with goat anti-PHA-L (1:2,000), rabbit anti-VGluT2 (1:2,000), and guinea pig anti-VGluT1 (1:20,000) primary antibodies diluted in the same solution (**Table 2**).

**TABLE 2.** Primary and secondary antibodies and dilutions for VGluT1, VGluT2 and *Phaseolus vulgaris* immunohistochemistry.

| ANTIBODY | IMMUNOGEN | MANUFACTURER | RRID | DILUTION |
| --- | --- | --- | --- | --- |
| ANTI-VGLUT1 | C-terminal residues 542–560 of rat VGluT1 | MilliporeSigma; guinea pig polyclonal; AB5905 | AB_2301751 | 1:20,000 |
| ANTI-VGLUT2 | C-terminal residues 510–582 of rat VGluT2 | Synaptic Systems; rabbit polyclonal; 135403 | AB_887883 | 1:2,000 |
| ANTI-PHASEOLUS VULGARIS AGGLUTININ (E+L) | Purified *Phaseolus vulgaris* agglutinin E+L | Vector Laboratories; goat polyclonal; AS-2224 | AB_10000080 | 1:2,000 |
| ALEXA FLUOR 647 DONKEY ANTI-GUINEA PIG IGG (H+L) | Guinea pig IgG (H+L) | Jackson ImmunoResearch; donkey polyclonal; 706-605-148 | AB_2340476 | 1:500 |
| ALEXA FLUOR 555 DONKEY ANTI-RABBIT IGG | Rabbit IgG | BioLegend; donkey polyclonal; 406412 | AB_2563181 | 1:500 |
| ALEXA FLUOR 488 DONKEY ANTI-GOAT IGG (H+L) | Goat IgG (H+L) | Jackson ImmunoResearch; donkey polyclonal; 705-545-147 | AB_2336933 | 1:1,000 |

After three washes in 0.1 M PBS, sections were incubated for 2 h with Alexa Fluor 488- conjugated donkey anti-goat IgG (H+L; 1:1,000), Alexa Fluor 555-conjugated donkey anti-rabbit IgG (1:500), and Alexa Fluor 647-conjugated donkey anti-guinea pig IgG (H+L; 1:500). Sections were then washed three times for 5 min in 0.1 M PBS, mounted onto gelatin-coated slides, and cover slipped with Eukitt quick-hardening mounting medium (Sigma-Aldrich, catalog no. 03989). Unless otherwise specified, all incubations were performed at room temperature. Control sections processed in parallel with the omission of the primary antibodies showed no detectable fluorescence under the acquisition settings used for the experimental sections.

### Image acquisition

#### Low-magnification imaging and anatomical delineation

All sections processed for PHA-L, VGluT1, and VGluT2 immunofluorescence were initially imaged using an Olympus BX51WI microscope equipped with a 10× UPlanApo objective (numerical aperture, 0.40) and a Zeiss AxioCam MRm Rev. 3.0 monochrome CCD camera (1388 × 1040 pixels; 6.45 × 6.45 µm pixel size; 12-bit digitization; Carl Zeiss, Germany) controlled by Neurolucida software (MBF Bioscience, Williston, VT, USA). Low-magnification tile scans were used to verify PHA-L injection sites, map the distribution of anterogradely labelled axons, and delineate cortical and subcortical regions of interest.

VGluT2 immunoreactivity was used as an anatomical marker to delineate cortical areas and layers, the dorsal lateral geniculate nucleus (dLGN), the lateral and medial subdivisions of the lateral posterior nucleus (LP_L_ and LP_M_), the superficial, intermediate, and deep layers of the superior colliculus (SC), the pontine nuclei (PN), and the striatum (ST). Anatomical structures were identified according to the mouse brain atlas of Paxinos and Franklin (2007).

#### Selection of sections and regions of interest

In each target structure, PHA-L-positive (PHA-L+) axons and boutons formed a projection field extending across multiple consecutive sections. Three approximately equidistant sections spanning the rostrocaudal extent of each projection field were selected for high-resolution analysis. For projections from V1 to the medial and lateral secondary visual areas, V2_M_ and V2_L_, respectively, six approximately equidistant sections were selected because of the relatively low density of labelled boutons in these pathways.

Within each selected cortical section, five regions of interest were sampled, corresponding to cortical layers 1, 2/3, 4, 5, and 6. In the lateral posterior nucleus, two regions of interest were sampled per section, corresponding to its lateral and medial subdivisions. In the superior colliculus, three regions of interest were sampled per section, corresponding to the superficial, intermediate, and deep layers. A single region of interest was sampled per section in each of the other subcortical targets, including the dLGN, PN, and ST.

Regions of interest were selected within PHA-L+ projection fields based on the distribution of anterogradely labelled axons. Within these regions, only PHA-L+ boutons displaying detectable VGluT1 and/or VGluT2 immunoreactivity were included in the quantitative analyses. PHA-L+ boutons without detectable labelling for VGluTs isoforms were not counted. Consequently, the reported proportions represent the relative prevalence of VGluT1 and VGluT2 among VGluT- immunoreactive PHA-L+ boutons, rather than the proportion of all PHA-L+ boutons expressing each transporter.

#### Confocal imaging

High-resolution image stacks were acquired using a Leica TCS SP8 confocal microscope equipped with a 63× HC PL APO CS2 oil-immersion objective (numerical aperture, 1.40). Alexa Fluor 488, Alexa Fluor 555, and Alexa Fluor 647 were excited sequentially using 488-, 552-, and 638-nm laser lines, respectively, to minimize spectral crosstalk. Images were acquired at 1024 × 1024 pixels using a zoom factor of 4 and sixfold line averaging.

Each stack comprised eight optical sections acquired at 0.3-µm intervals, corresponding to a sampled depth of 2.1 µm between the first and last optical planes and a voxel size of 45 × 45 × 300 nm. Because PHA-L immunolabelling extended throughout the section thickness whereas VGluT1 and VGluT2 antibody penetration was limited, confocal stacks were acquired close to the upper surface of each section.

### Data analysis

Confocal image stacks were imported into Fiji/ImageJ using the Bio-Formats plugin (Linkert et al. 2010; Schneider et al. 2012) The PHA-L, VGluT1, and VGluT2 channels were inspected both separately and as multichannel composites. Pixel intensities below 30 on an 8-bit scale of 0–255 were excluded from the analysis. This threshold was selected from control sections processed without the corresponding primary antibodies, in which nonspecific fluorescence did not exceed an intensity value of 30 under the acquisition conditions used (Landmann and Marbet 2004; Nakamura *et al*. 2007).

PHA-L+ axonal swellings were examined throughout each optical stack. Only PHA-L+ boutons containing detectable VGluT1 and/or VGluT2 immunoreactivity were included in the quantitative analysis, PHA-L+ boutons without detectable immunoreactivity for either transporter were not counted. Colocalization was assessed by examining the spatial overlap between PHA-L and VGluT immunoreactivity in individual optical planes and was confirmed using orthogonal projections. VGluT-positive (VGluT+) punctum was assigned to a PHA-L+ bouton only when the signals overlapped within the same optical planes and the VGluT+ punctum was contained within, or corresponded in size and position to, the PHA-L+ bouton.

PHA-L+ boutons and associated VGluT+ puncta were manually outlined at the optical plane in which each structure exhibited its largest cross-sectional profile. The maximal two-dimensional cross-sectional areas of both the PHA-L+ bouton and the associated VGluT+ punctum were measured in square micrometres. Bouton size was operationally defined as the maximal cross-sectional area of the PHA-L+ profile. The ratio of VGluT+ punctum area to PHA-L+ bouton area was subsequently calculated for each bouton. Structures intersecting the lateral boundaries of an image stack were excluded because their complete profiles could not be confirmed.

To facilitate systematic examination of regions containing dense axonal labelling, a 4 × 4 grid was superimposed on each image. Grid fields were examined sequentially, and each bouton was marked after measurement to prevent duplicate counting. Boutons were classified as VGluT1+, VGluT2+, or positive for both transporters according to the presence of suprathreshold immunoreactivity within the PHA-L+ profile.

Based on the Rayleigh criterion and the numerical aperture of the objective (Jonkman et al. 2003; see also Nakamura *et al*. 2007; Jonkman et al. 2020), the estimated lateral resolutions were 228, 253, and 290 nm for Alexa Fluor 488, 555, and 647, respectively, corresponding to theoretical circular profile areas of 0.041, 0.050, and 0.066 µm². These values were smaller than the smallest measured profiles in the corresponding channels, which were 0.061, 0.061, and 0.069 µm², respectively. These estimates were used to confirm that the smallest analyzed profiles exceeded the theoretical lateral resolution associated with their respective fluorescence channels.

For projection-level analyses, boutons were grouped according to their anatomical origin, target, and direction. Ascending projections comprised feedforward corticocortical (CC_A_), thalamocortical (TC), tectothalamic (TeT), and thalamostriatal (TS) pathways. Descending projections comprised feedback corticocortical (CC_D_), corticopontine (CP), corticostriatal (CS), corticothalamic (CT), corticotectal (CTe), and tectopontine (TeP) pathways. A small number of boutons were identified in lateral corticocortical projections between V2_L_ and V2_M_. Because only seven boutons were sampled from these lateral projections, they were described but excluded from inferential statistical analyses.

### Statistics

Statistical analyses were performed in R version 4.6.0 (R Core Team, 2026) using the lme4 (version 2.0.1), lmerTest (version 3.2.1), and emmeans (version 2.0.4) packages. The mouse was considered the experimental unit, and individual boutons were treated as repeated observations nested within confocal images and animals.

The prevalence of VGluT1+ and VGluT2+ boutons was summarized descriptively using counts and proportions according to projection direction and projection type. These proportions were calculated among PHA-L+ boutons in which either VGluT1 or VGluT2 was detected and therefore do not estimate the prevalence of VGluT expression among all PHA-L+ terminals. No inferential comparisons of isoform prevalence were performed because several projection types exhibited complete or near-complete separation of the two isoforms, with one isoform absent or represented by only a few boutons. Because VGluT1/VGluT2 double-labelled boutons occurred too infrequently to support reliable statistical inference, they were excluded from the statistical models. The seven boutons assigned to lateral corticocortical projections were retained in descriptive summaries but excluded from inferential analyses also because of their limited number.

Bouton area and VGluT-immunoreactive punctum area were analyzed separately using linear mixed-effect models fitted by maximum likelihood. Both outcomes were natural log transformed before analysis. VGluT isoform, projection direction, and their interaction were included as fixed effects, with random intercepts for animals and image nested within the animal. Model assumptions were evaluated using residual-versus-fitted plots and quantile–quantile plots of the residuals, and model singularity was assessed before inference. Both final models retained the animal and image random effects and showed no evidence of singularity.

Estimated marginal means and simple-effect contrasts were obtained using Satterthwaite-adjusted degrees of freedom. Contrasts compared VGluT1+ and VGluT2+ boutons within each projection direction and ascending and descending projections within each isoform. P values were adjusted across each predefined family of simple-effects contrasts using the Holm method. Estimated marginal means and 95% confidence intervals were back transformed from the logarithmic scale and are reported as geometric means on the original square-micrometre scale. All tests were two-sided, and adjusted P values below 0.05 were considered statistically significant. No prospective sample-size calculation was performed because the analyses were conducted on an existing anatomical dataset.

## Results

### Regional distribution of VGluT1 and VGluT2 immunoreactivity

VGluT1 and VGluT2 exhibited distinct regional and laminar distributions throughout the cortical and subcortical structures examined (**Figure 2**). In the neocortex, VGluT1 immunoreactivity was detected in all cortical layers. Labelling was particularly prominent in layer 1 and progressively decreased across layers 2 and 3, whereas layer 4 generally exhibited weaker immunoreactivity than the surrounding layers. A similar laminar pattern was observed in parietal, temporal, and occipital visual areas (**Figure 2A, C**).

**Figure 2.**
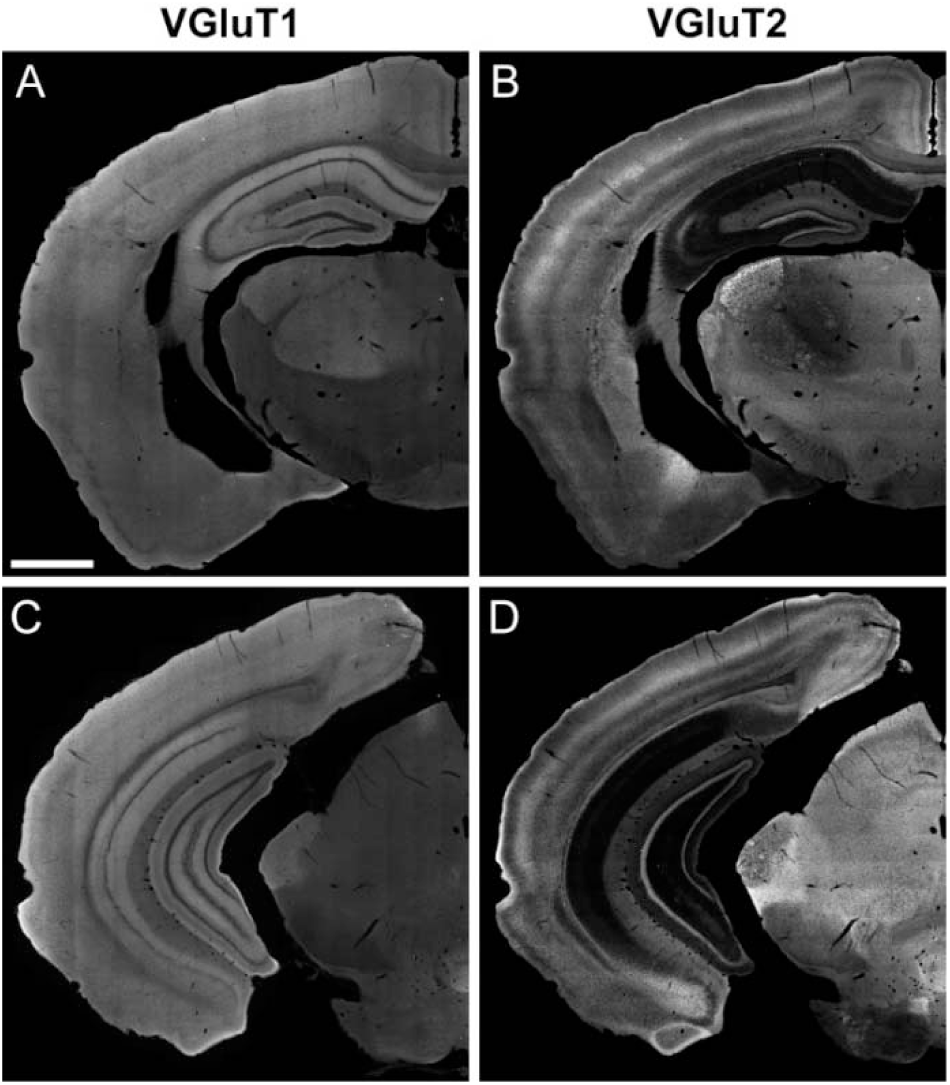
Regional and laminar distributions of VGluT1 and VGluT2 immunoreactivity in the mouse brain. Representative low-magnification fluorescence images of coronal brain sections acquired through the VGluT1 (A, C) and VGluT2 (B, D) channels. Panels A and B show the same rostral section, approximately 1.8 mm posterior to bregma, whereas panels C and D show the same more caudal section, approximately 2.8 mm posterior to bregma. (**A-B**) Scale bar = 1 mm.

In subcortical structures, moderate VGluT1 immunoreactivity was observed in the dorsal lateral geniculate nucleus, the ventral posteromedial and ventral posterolateral thalamic nuclei, the medial geniculate complex, the posterior thalamic nucleus, and the lateral posterior thalamic nucleus. By contrast, little or no detectable VGluT1 immunoreactivity was observed in the ventral lateral geniculate nucleus, medial thalamic nuclei, hypothalamus, or zona incerta. Weak VGluT1 labelling was also present in the superficial layers of the superior colliculus (**Figure 2A, C**).

VGluT2 immunoreactivity displayed a complementary cortical pattern. Labelling was prominent in layer 1, weak in layers 2/3, and particularly intense in layer 4 throughout the cortical mantle. Labelling was weaker in layer 5 and moderate in the superficial portion of layer 6a (**Figure 2B, D**).

Strong VGluT2 immunoreactivity was observed in the dorsal lateral geniculate nucleus, particularly within its shell region, whereas labelling in its core was moderate. The ventral lateral geniculate nucleus also displayed VGluT2 immunoreactivity. VGluT2+ puncta were evident in the lateral posterior nucleus, the ventral posteromedial and ventral posterolateral nuclei, and several subdivisions of the medial geniculate complex. Strong VGluT2 immunoreactivity was also observed in the optic and superficial grey layers of the superior colliculus, whereas its intermediate and deep layers exhibited more moderate labelling (**Figure 2B, D**).

### Anatomical identification and sampling of VGluT-positive boutons

PHA-L injections successfully targeted V1, V2_L_, dLGN, LP, or SC (**Figure 1**). The stereotaxic coordinates, injection duration, and individual cases retained for each injection site are presented in **Table 1**. Only animals in which the injection core remained confined to the intended structure were included. PHA-L-labelled axons and boutons were subsequently examined in cortical and subcortical targets of each injected structure. Representative examples of PHA-L+ boutons assessed for VGluT1 and VGluT2 immunoreactivity are shown separately in **Figures 3 and 4**, respectively, because the relative abundance of the two transporter isoforms differed markedly among pathways.

**Figure 3.**
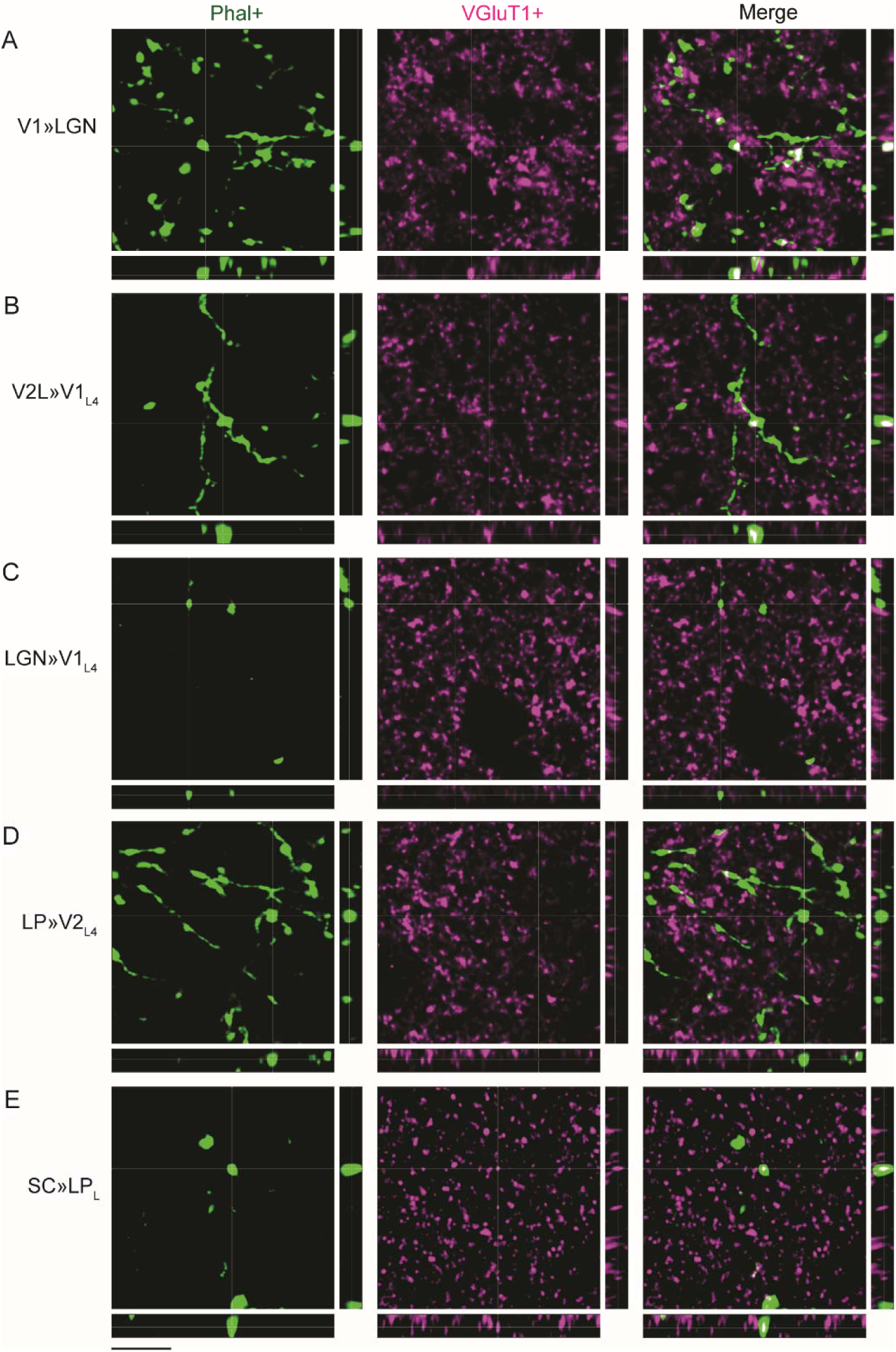
Identification of VGluT1 immunoreactivity in PHA-L-labelled boutons from anatomically identified visual pathways. Representative confocal image stacks are presented for the projections indicated in rows **A–E**. PHA-L-labelled axons and boutons are shown in green and VGluT1 immunoreactivity in magenta. The large panels correspond to the xy plane, whereas the narrow panels below and to the right show the corresponding xz and yz orthogonal views. A PHA-L-labelled bouton was considered VGluT1+ when spatial overlap between the two signals was confirmed in all three orthogonal planes. (**A-E**) Scale bar = 3 µm.

**Figure 4.**
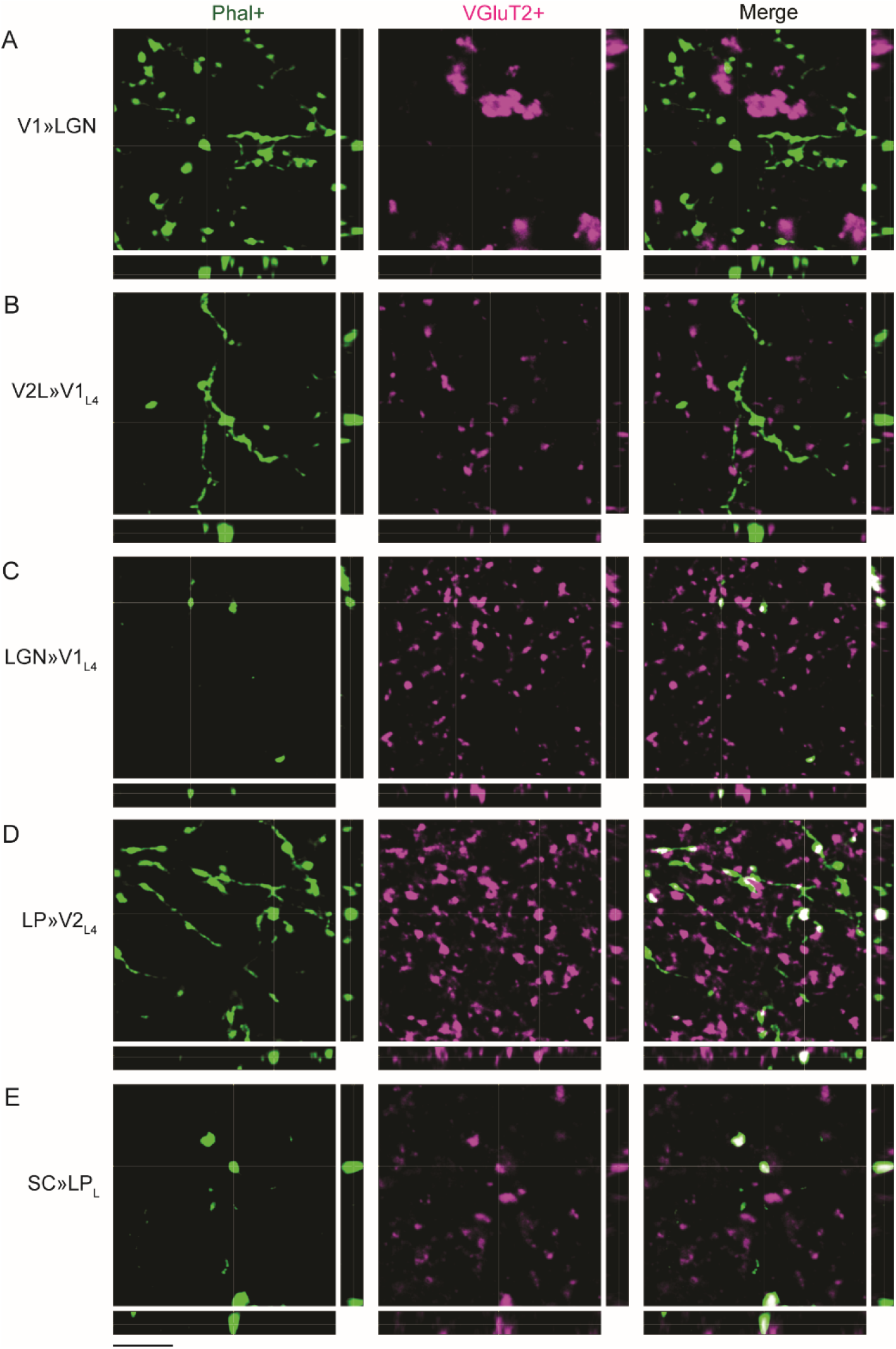
Identification of VGluT2 immunoreactivity in PHA-L-labelled boutons from anatomically identified visual pathways. Representative confocal image stacks are presented for the projections indicated in rows A–E. PHA-L-labelled axons and boutons are shown in green and VGluT2 immunoreactivity in magenta. The large panels correspond to the xy plane, whereas the narrow panels below and to the right show the corresponding xz and yz orthogonal views. A PHA-L-labelled bouton was considered VGluT2+ when spatial overlap between the two signals was confirmed in all three orthogonal planes. (**A-E**) Scale bar = 3 µm.

The quantitative dataset comprised 2,178 PHA-L-labelled boutons classified as singly positive for either VGluT1 or VGluT2. Cortical and subcortical counts are reported in **Tables 3 and 4**, respectively. Among these singly labelled boutons, 1,238 (56.8%) were VGluT1+ and 940 (43.2%) were VGluT2+. An additional 12 PHA-L-labelled boutons exhibited detectable immunoreactivity for both VGluT1 and VGluT2, yielding a total of 2,190 VGluT-immunoreactive boutons examined. Dual-labelled boutons therefore represented approximately 0.55% of the total sample.

**TABLE 3.**
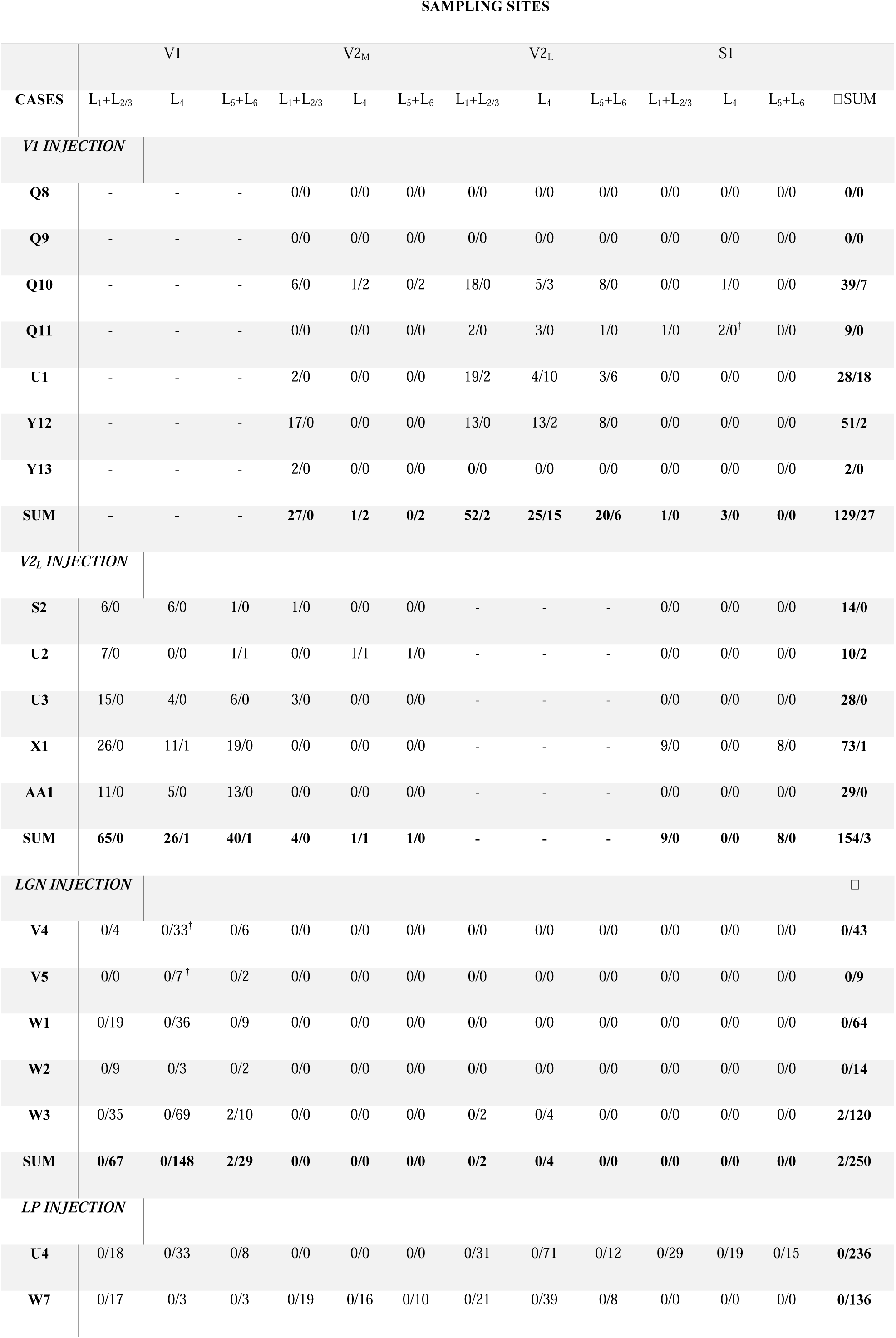
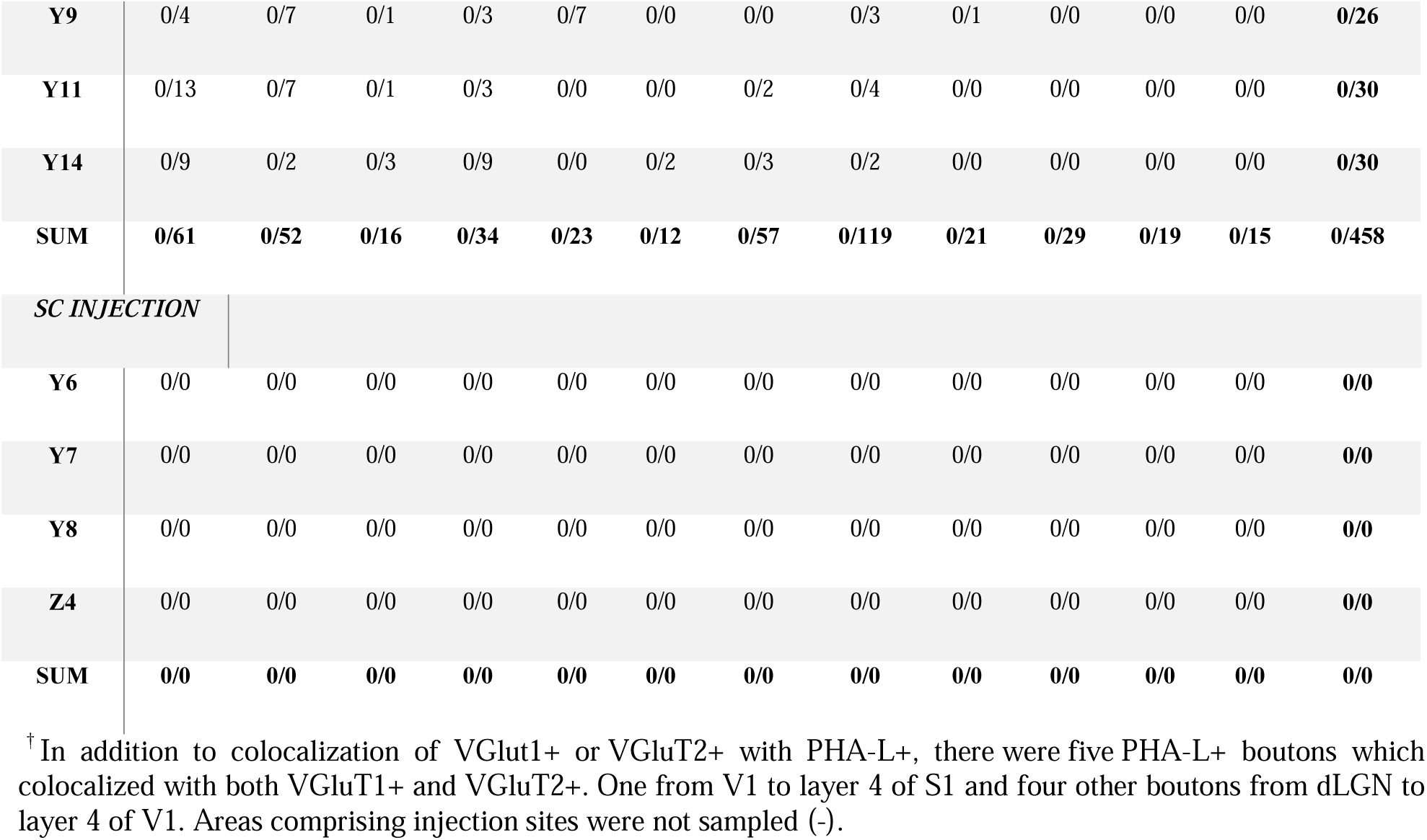
Total number of sampled VGluT1+/ VGluT2+ boutons colocalized with PHA-L+ in cortical areas V1, medial (V2_M_) and lateral (V2_L_) extrastriate visual and primary somatosensory cortices (S1) in supragranular (L_1_+L_2/3_), granular (L_4_) and infragranular (L_5_+L _6_) layers.

| SAMPLING SITES |  |  |  |  |  |  |  |  |  |  |  |  |  |
| --- | --- | --- | --- | --- | --- | --- | --- | --- | --- | --- | --- | --- | --- |
|  | V1 |  |  | V2<sub>M</sub> |  |  | V2<sub>L</sub> |  |  | S1 |  |  |  |
| CASES | L<sub>1</sub>+L<sub>2/3</sub> | L<sub>4</sub> | L<sub>5</sub>+L<sub>6</sub> | L<sub>1</sub>+L<sub>2/3</sub> | L<sub>4</sub> | L<sub>5</sub>+L<sub>6</sub> | L<sub>1</sub>+L<sub>2/3</sub> | L<sub>4</sub> | L<sub>5</sub>+L<sub>6</sub> | L<sub>1</sub>+L<sub>2/3</sub> | L<sub>4</sub> | L<sub>5</sub>+L<sub>6</sub> | □SUM |
| **V1 INJECTION** |  |  |  |  |  |  |  |  |  |  |  |  |  |
| **Q8** | - | - | - | 0/0 | 0/0 | 0/0 | 0/0 | 0/0 | 0/0 | 0/0 | 0/0 | 0/0 | **0/0** |
| **Q9** | - | - | - | 0/0 | 0/0 | 0/0 | 0/0 | 0/0 | 0/0 | 0/0 | 0/0 | 0/0 | **0/0** |
| **Q10** | - | - | - | 6/0 | 1/2 | 0/2 | 18/0 | 5/3 | 8/0 | 0/0 | 1/0 | 0/0 | **39/7** |
| **Q11** | - | - | - | 0/0 | 0/0 | 0/0 | 2/0 | 3/0 | 1/0 | 1/0 | 2/0<sup>†</sup> | 0/0 | **9/0** |
| **U1** | - | - | - | 2/0 | 0/0 | 0/0 | 19/2 | 4/10 | 3/6 | 0/0 | 0/0 | 0/0 | **28/18** |
| **Y12** | - | - | - | 17/0 | 0/0 | 0/0 | 13/0 | 13/2 | 8/0 | 0/0 | 0/0 | 0/0 | **51/2** |
| **Y13** | - | - | - | 2/0 | 0/0 | 0/0 | 0/0 | 0/0 | 0/0 | 0/0 | 0/0 | 0/0 | **2/0** |
| **SUM** | - | - | - | **27/0** | **1/2** | **0/2** | **52/2** | **25/15** | **20/6** | **1/0** | **3/0** | **0/0** | **129/27** |
| **V2<sub>L</sub> INJECTION** |  |  |  |  |  |  |  |  |  |  |  |  |  |
| **S2** | 6/0 | 6/0 | 1/0 | 1/0 | 0/0 | 0/0 | - | - | - | 0/0 | 0/0 | 0/0 | **14/0** |
| **U2** | 7/0 | 0/0 | 1/1 | 0/0 | 1/1 | 1/0 | - | - | - | 0/0 | 0/0 | 0/0 | **10/2** |
| **U3** | 15/0 | 4/0 | 6/0 | 3/0 | 0/0 | 0/0 | - | - | - | 0/0 | 0/0 | 0/0 | **28/0** |
| **X1** | 26/0 | 11/1 | 19/0 | 0/0 | 0/0 | 0/0 | - | - | - | 9/0 | 0/0 | 8/0 | **73/1** |
| **AA1** | 11/0 | 5/0 | 13/0 | 0/0 | 0/0 | 0/0 | - | - | - | 0/0 | 0/0 | 0/0 | **29/0** |
| **SUM** | **65/0** | **26/1** | **40/1** | **4/0** | **1/1** | **1/0** | - | - | - | **9/0** | **0/0** | **8/0** | **154/3** |
| **LGN INJECTION** |  |  |  |  |  |  |  |  |  |  |  |  |  |
| **V4** | 0/4 | 0/33<sup>†</sup> | 0/6 | 0/0 | 0/0 | 0/0 | 0/0 | 0/0 | 0/0 | 0/0 | 0/0 | 0/0 | **0/43** |
| **V5** | 0/0 | 0/7<sup>†</sup> | 0/2 | 0/0 | 0/0 | 0/0 | 0/0 | 0/0 | 0/0 | 0/0 | 0/0 | 0/0 | **0/9** |
| **W1** | 0/19 | 0/36 | 0/9 | 0/0 | 0/0 | 0/0 | 0/0 | 0/0 | 0/0 | 0/0 | 0/0 | 0/0 | **0/64** |
| **W2** | 0/9 | 0/3 | 0/2 | 0/0 | 0/0 | 0/0 | 0/0 | 0/0 | 0/0 | 0/0 | 0/0 | 0/0 | **0/14** |
| **W3** | 0/35 | 0/69 | 2/10 | 0/0 | 0/0 | 0/0 | 0/2 | 0/4 | 0/0 | 0/0 | 0/0 | 0/0 | **2/120** |
| **SUM** | **0/67** | **0/148** | **2/29** | **0/0** | **0/0** | **0/0** | **0/2** | **0/4** | **0/0** | **0/0** | **0/0** | **0/0** | **2/250** |
| **LP INJECTION** |  |  |  |  |  |  |  |  |  |  |  |  |  |
| **U4** | 0/18 | 0/33 | 0/8 | 0/0 | 0/0 | 0/0 | 0/31 | 0/71 | 0/12 | 0/29 | 0/19 | 0/15 | **0/236** |
| **W7** | 0/17 | 0/3 | 0/3 | 0/19 | 0/16 | 0/10 | 0/21 | 0/39 | 0/8 | 0/0 | 0/0 | 0/0 | **0/136** |
| **Y9** | 0/4 | 0/7 | 0/1 | 0/3 | 0/7 | 0/0 | 0/0 | 0/3 | 0/1 | 0/0 | 0/0 | 0/0 | **0/26** |
| **Y11** | 0/13 | 0/7 | 0/1 | 0/3 | 0/0 | 0/0 | 0/2 | 0/4 | 0/0 | 0/0 | 0/0 | 0/0 | **0/30** |
| **Y14** | 0/9 | 0/2 | 0/3 | 0/9 | 0/0 | 0/2 | 0/3 | 0/2 | 0/0 | 0/0 | 0/0 | 0/0 | **0/30** |
| **SUM** | **0/61** | **0/52** | **0/16** | **0/34** | **0/23** | **0/12** | **0/57** | **0/119** | **0/21** | **0/29** | **0/19** | **0/15** | **0/458** |
| **SC INJECTION** |  |  |  |  |  |  |  |  |  |  |  |  |  |
| **Y6** | 0/0 | 0/0 | 0/0 | 0/0 | 0/0 | 0/0 | 0/0 | 0/0 | 0/0 | 0/0 | 0/0 | 0/0 | **0/0** |
| **Y7** | 0/0 | 0/0 | 0/0 | 0/0 | 0/0 | 0/0 | 0/0 | 0/0 | 0/0 | 0/0 | 0/0 | 0/0 | **0/0** |
| **Y8** | 0/0 | 0/0 | 0/0 | 0/0 | 0/0 | 0/0 | 0/0 | 0/0 | 0/0 | 0/0 | 0/0 | 0/0 | **0/0** |
| **Z4** | 0/0 | 0/0 | 0/0 | 0/0 | 0/0 | 0/0 | 0/0 | 0/0 | 0/0 | 0/0 | 0/0 | 0/0 | **0/0** |
| **SUM** | **0/0** | **0/0** | **0/0** | **0/0** | **0/0** | **0/0** | **0/0** | **0/0** | **0/0** | **0/0** | **0/0** | **0/0** | **0/0** |

<sup>†</sup> In addition to colocalization of VGluT1+ or VGluT2+ with PHA-L+, there were five PHA-L+ boutons which colocalized with both VGluT1+ and VGluT2+. One from V1 to layer 4 of S1 and four other boutons from dLGN to layer 4 of V1. Areas comprising injection sites were not sampled (-).

**TABLE 4.** Total number of sampled VGluT1+/ VGluT2+ boutons colocalized with PHA-L+ in subcortical areas.

| SAMPLING SITES |  |  |  |  |  |  |  |  |  |
| --- | --- | --- | --- | --- | --- | --- | --- | --- | --- |
| CASES | dLGN | LPM | LPL | SCS | SCI | SCD | ST | PN | SUM |
| **V1 INJECTION** |  |  |  |  |  |  |  |  |  |
| Q8 | 9/0 | 0/0 | 1/0 | 4/0 | 0/0 | 0/0 | 0/0 | 0/0 | 14/0 |
| Q9 | 15/0 | 3/0 | 0/0 | 14/0 | 0/0 | 1/0 | 0/0 | 5/0 | 38/0 |
| Q10 | 263/0 | 2/0 | 19/0 | 8/0 | 1/0 | 0/0 | 5/0 | 0/0 | 298/0 |
| Q11 | 40/0 | 2/0 | 5/0 | 8/0 | 0/0 | 0/0 | 6/0 | 3/0 | 64/0 |
| U1 | 88/0 | 0/0 | 3/0 | 5/0 | 6/1 | 0/0 | 17/1<sup>†</sup> | 0/0 | 119/1 |
| Y12 | 19/0 | 7/0 | 4/0 | 12/0 | 0/0 | 0/0 | 3/0 | 12/0 | 57/0 |
| Y13 | 10/0 | 0/0 | 0/0 | 5/1 | 1/0 | 0/0 | 0/0 | 1/0 | 17/1 |
| SUM | 444/0 | 14/0 | 32/0 | 56/1 | 8/1 | 1/0 | 31/1 | 21/0 | 607/2 |
| **V2<sub>L</sub> INJECTION** |  |  |  |  |  |  |  |  |  |
| S2 | 0/0 | 0/0 | 18/0 | 4/0 | 4/0 | 0/0 | 2/0 | 3/0 | 31/0 |
| U2 | 1/0 | 0/0 | 47/0 | 0/0 | 0/0 | 0/0 | 17/1 | 11/0 | 76/1 |
| U3 | 27/0 | 0/0 | 15/0 | 8/0 | 5/0 | 4/0 | 10/2 | 5/0 | 74/2 |
| X1 | 0/0 | 3/0 | 55/0 | 0/0 | 5/0 | 0/0 | 21/0 | 15/0 | 99/0 |
| AA1 | 8/0 | 4/0 | 19/0 | 9/1 | 4/0 | 4/0 | 13/0 | 4/0 | 65/1 |
| SUM | 36/0 | 7/0 | 154/0 | 21/1 | 18/0 | 8/0 | 63/3 | 38/0 | 345/4 |
| **LGN INJECTION** |  |  |  |  |  |  |  |  |  |
| V4 | - | 0/0 | 0/0 | 0/0 | 0/0 | 0/0 | 0/0 | 0/0 | 0/0 |
| V5 | - | 0/0 | 0/0 | 0/0 | 0/0 | 0/0 | 0/0 | 0/0 | 0/0 |
| W1 | - | 0/0 | 0/0 | 0/0 | 0/0 | 0/0 | 0/0 | 0/0 | 0/0 |
| W2 | - | 0/0 | 0/0 | 0/0 | 0/0 | 0/0 | 0/0 | 0/0 | 0/0 |
| W3 | - | 0/0 | 0/0 | 0/0 | 0/0 | 0/0 | 0/0 | 0/0 | 0/0 |
| SUM | - | 0/0 | 0/0 | 0/0 | 0/0 | 0/0 | 0/0 | 0/0 | 0/0 |
| **LP INJECTION** |  |  |  |  |  |  |  |  |  |
| U4 | 0/0 | - | - | 0/0 | 0/0 | 0/0 | 0/1 | 0/0 | 0/1 |
| W7 | 0/0 | - | - | 0/0 | 0/0 | 0/0 | 0/4 | 0/0 | 0/4 |
| Y9 | 0/0 | - | - | 0/0 | 0/0 | 0/0 | 0/1 | 0/0 | 0/1 |
| Y11 | 0/0 | - | - | 0/0 | 0/0 | 0/0 | 0/0 | 0/0 | 0/0 |
| Y14 | 0/0 | - | - | 0/0 | 0/0 | 0/0 | 0/0 | 0/0 | 0/0 |
| SUM | 0/0 | - | - | 0/0 | 0/0 | 0/0 | 0/6 | 0/0 | 0/6 |
| **SC INJECTION** |  |  |  |  |  |  |  |  |  |
| Y6 | 1/9<sup>†</sup> | 0/16 | 0/15 | - | - | - | 0/0 | 0/13 | 1/53 |
| Y7 | 0/0 | 0/27 | 0/22 | - | - | - | 0/0 | 0/6 | 0/55 |
| Y8 | 0/27 | 0/15 | 0/12<sup>†</sup> | - | - | - | 0/0 | 0/6 | 0/60 |
| Z4 | 0/2 | 0/8 | 0/8<sup>†</sup> | - | - | - | 0/0 | 0/4 | 0/22 |
| SUM | 1/38 | 0/66 | 0/57 | - | - | - | 0/0 | 0/29 | □1/190 |

<sup>†</sup> In addition to colocalization of VGluT1+ or VGluT2+ with PHA-L+, there were seven PHA-L+ boutons within this number colocalized with both VGluT1+ and VGluT2+. One bouton from V1 to ST and six more boutons from SC were observed with three to dLGN and three to LPL. Areas comprising injection sites were not sampled (-).

The singly labelled population was dominated by thalamocortical and corticothalamic projections, which comprised 710 and 688 boutons, respectively. Corticocortical projections accounted for an additional 313 boutons.

### VGluT phenotype differs across anatomically identified pathways

The distribution of VGluT isoforms differed markedly among projection classes (**Figure 5A**; **Tables 3 and 4**). Corticothalamic and corticopontine boutons were exclusively classified as VGluT1+ in the singly labelled dataset. VGluT1 also predominated in descending corticocortical, corticotectal, and corticostriatal projections, accounting for 98.5%, 97.4%, and 96.9% of the singly labelled boutons sampled from these pathways, respectively. In contrast, all singly labelled tectopontine boutons were VGluT2+.

**Figure 5.**
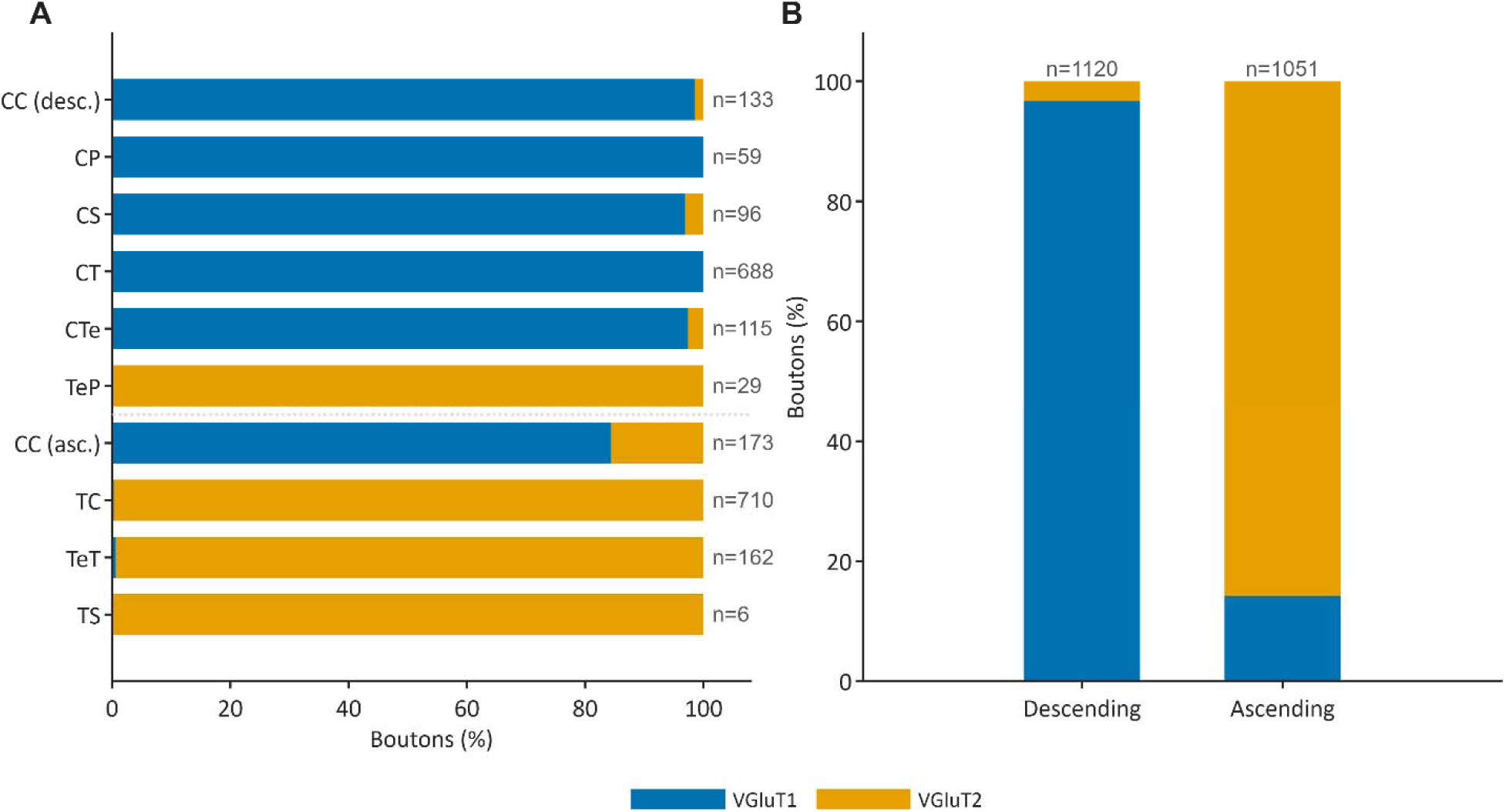
VGluT phenotype differs among anatomically identified projection classes. (**A**) Relative proportions of singly labelled PHA-L+ boutons classified as VGluT1+ or VGluT2+ within each projection class. The dashed horizontal line separates descending from ascending projection classes. (**B**) Distribution of VGluT1+ and VGluT2+ boutons after pooling all descending or ascending projection classes. Values above or beside the bars indicate the total number of singly labelled boutons included in each category.

Ascending thalamocortical and tectothalamic projections were characterized by an almost exclusive expression of VGluT2. Among the 710 singly labelled thalamocortical boutons analysed, 708 (99.7%) were VGluT2+ (**Table 3**). Similarly, 161 of 162 singly labelled tectothalamic boutons (99.4%) were VGluT2+ (**Table 4**). All six boutons sampled from the LP-to-striatum thalamostriatal projection were also VGluT2+.

The transporter distribution was less homogeneous in ascending corticocortical pathways. Of the 173 singly labelled boutons assigned to this projection class, 146 (84.4%) were VGluT1+ and 27 (15.6%) were VGluT2+ (**Table 3**). Thus, unlike the other ascending pathways examined, ascending corticocortical projections remained predominantly associated with VGluT1.

Following injections in V1, 156 singly labelled boutons were sampled in V2_M_, V2_L_, and S1 (**Table 3**). Of these, 129 (82.7%) were VGluT1+ and 27 (17.3%) were VGluT2+. VGluT2+ boutons were concentrated primarily in the projections to V2_L_, particularly in layer 4. An additional bouton in the V1-to-S1 projection exhibited VGluT1/VGluT2 colocalization and was in layer 4. Among the 609 singly labelled boutons sampled in subcortical targets following V1 injections, 607 were VGluT1+ and two were VGluT2+ (**Table 4**). One additional double-labelled bouton was observed in the V1-to-striatum projection.

Following injections in V2_L_, 157 singly labelled corticocortical boutons were sampled in V1, V2M, and S1 (**Table 3**). Of the 133 boutons sampled in the feedback projection to V1, 131 were VGluT1+ and two were VGluT2+. All 17 boutons examined in the V2_L_-to-S1 projection were VGluT1+. Among the 349 singly labelled boutons sampled in subcortical targets, 345 were VGluT1+ and four were VGluT2+ (**Table 4**). Thus, both corticocortical feedback and corticofugal projections originating in V2L were strongly dominated by VGluT1.

In contrast, projections originating in the dLGN and LP were almost exclusively VGluT2+. Following dLGN injections, 250 of 252 singly labelled cortical boutons were VGluT2+ (**Table 3**). In the dLGN-to-V1 projection, VGluT2+ boutons were most abundant in layer 4, although they were also detected in supra- and infragranular layers. Four additional boutons in layer 4 of V1 exhibited colocalization of VGluT1 and VGluT2. All 458 cortical boutons sampled following LP injections were singly VGluT2+ and were distributed across multiple cortical layers, with substantial numbers in layers 1 and 4 (**Table 3**). All six boutons sampled in the LP-to-striatum projection were also VGluT2+ (**Table 4**).

Projections originating in the SC were similarly dominated by VGluT2. Of the 162 singly labelled tectothalamic boutons sampled in the dLGN and LP, 161 were VGluT2+, whereas all 29 singly labelled tectopontine boutons were VGluT2+ (**Table 4**). Six additional tectothalamic boutons exhibited VGluT1/VGluT2 colocalization, including three boutons in the SC-to-dLGN projection and three in the SC-to-LP projection.

### VGluT distribution according to projection direction

When singly labelled boutons were grouped according to anatomical direction, VGluT1 and VGluT2 exhibited largely complementary distributions (**Figure 5B**; **Tables 3 and 4**). Among 1,120 boutons assigned to descending projections, 1,083 (96.7%) were VGluT1+ and 37 (3.3%) were VGluT2+. Conversely, among 1,051 boutons assigned to ascending projections, 902 (85.8%) were VGluT2+ and 149 (14.2%) were VGluT1+.

The predominance of VGluT2 among ascending projections was attributable primarily to thalamocortical and tectothalamic pathways. Ascending corticocortical projections represented an important exception because they remained predominantly VGluT1+. Likewise, the tectopontine pathway constituted an exception among descending projections because all the singly labelled boutons therein was VGluT2+. The seven singly labelled boutons assigned to a lateral corticocortical projection were not included in the direction-based summaries; six were VGluT1+ and one was VGluT2+.

### VGluT1 and VGluT2 colocalization is rare and pathway-specific

Twelve PHA-L-labelled boutons exhibited detectable immunoreactivity for both VGluT1 and VGluT2, corresponding to approximately 0.55% of all VGluT-immunoreactive boutons examined (**Tables 3 and 4**). Five double-labelled boutons were identified in cortical targets: one in layer 4 of S1 following a V1 injection and four in layer 4 of V1 following dLGN injections. Seven additional double-labelled boutons were identified in subcortical targets: one in the striatum following a V1 injection and six in tectothalamic projections originating in the SC, including three in the dLGN and three in LP.

Dual labelling was therefore rare and predominantly observed in ascending thalamocortical or tectothalamic pathways. Given the small number of double-labelled boutons and their distinct molecular phenotype, morphometric analyses were restricted to boutons unambiguously classified as singly VGluT1+ or VGluT2+. Dual-labelled boutons were consequently reported descriptively but were not included in the mixed-effect analyses.

### Bouton area depends on VGluT isoform and projection direction

Morphometric analyses included 2,171 singly labelled boutons from 26 animals and 472 animals–image combinations. The seven boutons assigned to lateral corticocortical projections were excluded because the inferential models compared ascending and descending projections only.

Bouton area showed a significant interaction between VGluT isoform and projection direction (β = 0.450, SE = 0.142, *t*(550.70) = 3.17, *p* = 0.0016; **Figure 6A**). The estimated geometric mean area of VGluT1+ boutons was 0.613 µm² (95% CI: 0.534–0.705 µm²) in descending projections and 0.387 µm² (95% CI: 0.326–0.459 µm²) in ascending projections. VGluT1+ boutons were therefore larger in descending than in ascending projections (Holm-adjusted *p* < 0.001).

**Figure 6.**
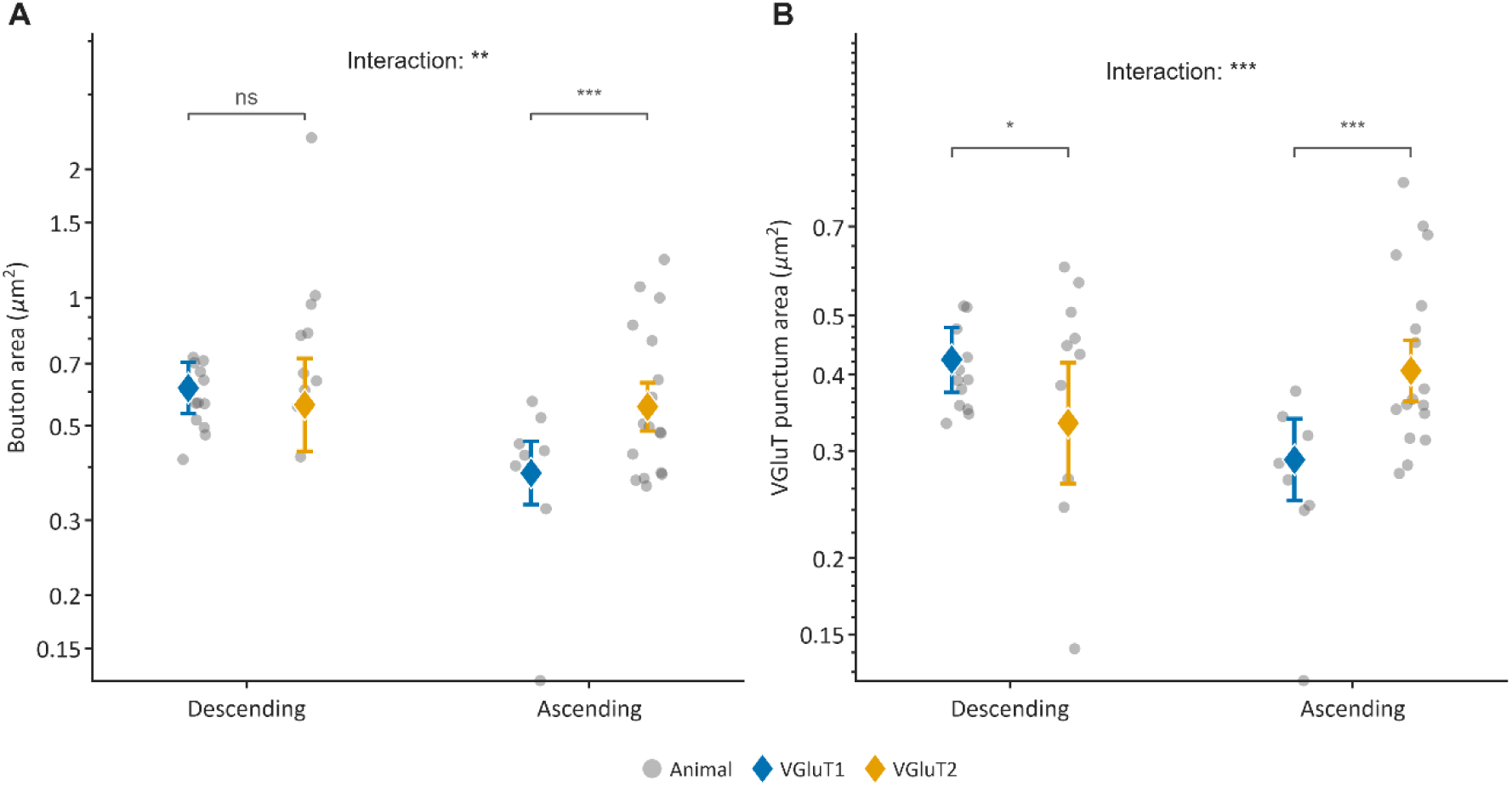
Bouton and VGluT-positive punctum areas vary according to VGluT isoform and projection direction. **(A)** Area of PHA-L-labelled boutons and **(B)** area of the corresponding VGluT-positive puncta in descending and ascending projections. Grey circles represent animal-level values. Coloured diamonds and error bars indicate back-transformed model-estimated geometric means and their 95% confidence intervals for VGluT1+ (blue) and VGluT2+ (orange) boutons. Statistical brackets show Holm-adjusted pairwise comparisons between VGluT isoforms within each projection direction. Interaction significance was obtained from the VGluT isoform × projection direction term of the mixed-effect model. Analyses included singly labelled boutons from ascending and descending projections; lateral corticocortical and double-labelled boutons were excluded. The ordinate is displayed on a logarithmic scale. ns, p ≥ 0.05; *p < 0.05; **p < 0.01; ***p < 0.001.

For VGluT2+ boutons, the estimated geometric mean area was 0.560 µm² (95% CI: 0.436–0.719 µm²) in descending projections and 0.554 µm² (95% CI: 0.486–0.631 µm²) in ascending projections. Bouton area did not differ significantly between descending and ascending VGluT2+ projections (Holm-adjusted *p* = 0.935).

Within descending projections, VGluT1+ and VGluT2+ boutons did not differ significantly in area (Holm-adjusted *p* = 0.485). Within ascending projections, however, VGluT2+ boutons were larger than VGluT1+ boutons (Holm-adjusted *p* < 0.001). Thus, the predicted association between VGluT2 and larger boutons was supported specifically within ascending projections rather than across all pathways.

### VGluT-positive punctum area depends on transporter isoform and projection direction

VGluT-positive punctum area also showed a significant interaction between transporter isoform and projection direction (β = 0.575, SE = 0.130, *t*(545.49) = 4.44, *p* < 0.001; **Figure 6B**). For VGluT1, the estimated geometric mean punctum area was 0.423 µm² (95% CI: 0.374–0.478 µm²) in descending projections and 0.290 µm² (95% CI: 0.249–0.338 µm²) in ascending projections. VGluT1+ puncta were therefore larger in descending than in ascending pathways (Holm-adjusted *p* < 0.001).

For VGluT2, the estimated geometric mean punctum area was 0.333 µm² (95% CI: 0.265–0.419 µm²) in descending projections and 0.406 µm² (95% CI: 0.361–0.456 µm²) in ascending projections. This directional difference did not reach statistical significance after correction for multiple comparisons (Holm-adjusted *p* = 0.086).

Within descending projections, VGluT1+ puncta were modestly larger than VGluT2+ puncta (Holm-adjusted *p* = 0.047). The opposite pattern was observed in ascending projections, in which VGluT2+ puncta were larger than VGluT1+ puncta (Holm-adjusted *p* < 0.001). The association between transporter isoform and terminal morphology therefore depended on projection direction.

## Discussion

### General organization of VGluT phenotypes across visual pathways

The present study provides a projection-specific characterization of VGluT1 and VGluT2 expression across multiple components of the mouse visual system. By combining anterograde tracing with simultaneous detection of both isoforms, we directly identified the VGluT phenotype of boutons arising from cortical, thalamic, and tectal structures. The results reveal a broad, but non-absolute, descending–ascending organization. Among singly labelled VGluT+ boutons, corticothalamic and corticopontine projections were exclusively classified as VGluT1+, while corticocortical feedback, corticostriatal, and corticotectal pathways were strongly dominated by VGluT1. Conversely, thalamocortical, tectothalamic, thalamostriatal, and tectopontine pathways were dominated by VGluT2. This organization is consistent with comparative studies associating VGluT1 with intrinsic cortical and descending corticofugal pathways and VGluT2 with subcortical and thalamocortical projections (Kaneko and Fujiyama 2002; Balaram, Hackett*, et al.* 2011; Balaram *et al*. 2013; Balaram *et al*. 2015; Abbas Farishta *et al*. 2022). The present findings extend this regional and laminar evidence by demonstrating transporter phenotype directly within boutons belonging to anatomically identified pathways.

Projection direction nevertheless provides an incomplete account of transporter phenotype. The principal exception was the ascending corticocortical projections, which expressed predominantly VGluT1. VGluT2+ boutons were mainly in layer 4 of projection from V1 to V2_L_. This heterogeneity indicates that corticocortical feedforward projections are not molecularly equivalent to ascending thalamocortical pathways and may comprise presynaptic populations differing in laminar origin, termination patterns, transporter phenotype, or physiological properties. The tectopontine pathway represented the converse exception: although anatomically descending, all singly labelled boutons sampled from this pathway were VGluT2+. Together, these exceptions suggest that neuronal origin and pathway identity constrain VGluT phenotype more strongly than anatomical direction alone.

### Corticocortical projections reveal molecular heterogeneity within the feedforward–feedback hierarchy

VGluT1 was the most prevalent isoform in both feedforward and feedback corticocortical projections. VGluT2 was present in the feedforward projections from V1 to the extrastriate visual areas V2_L_ and V2_M_ and was detected only rarely in the feedback projections from V2_L_-to-V1. Thus, although corticocortical projections were predominantly associated with VGluT1 regardless of direction, feedforward projections exhibited greater molecular heterogeneity than feedback projections.

The laminar distribution of the VGluT+ boutons was broadly consistent with the organization of corticocortical pathways described in rodents. Feedforward corticocortical projections target all cortical layers, including layer 4, whereas feedback projections preferentially terminate in supra- and infragranular layers and generally provide less prominent innervation of layer 4 (Coogan and Burkhalter 1990, 1993). A comparable organization having been described in primate visual cortex (Rockland and Pandya 1979). In the present study, VGluT2+ boutons in the V1-to-V2_L_ projection were concentrated primarily in layer 4, whereas only a few were detected in the V2_L_- to-V1 feedback projection. VGluT1+ boutons were distributed more broadly across the cortical depth. In the V1-to-V2_L_ feedforward projection, they were abundant in the supra- and infragranular layers but were also present in layer 4. In the V2_L_-to-V1 feedback projection, they predominated in the supra- and infragranular layers, although they were detected across all cortical layers. Layer 4 innervation was therefore not exclusively associated with VGluT2 expression.

These observations complement previous morphological evidence showing that feedforward corticocortical axons from V1 form numerous *en passant* boutons in layer 4, despite exhibiting limited axonal branching within this layer (Massé et al. 2016). The present results show that these layer 4 boutons are molecularly heterogeneous and may express either VGluT1 or VGluT2. Consequently, the feedforward character and laminar termination pattern of a corticocortical projection do not uniquely determine its VGluT phenotype. This result distinguishes ascending corticocortical projections from the nearly uniform VGluT2 phenotype observed in the ascending thalamocortical pathways examined here.

The cellular origin of this molecular heterogeneity cannot be established from the present terminal-based analysis. Although the cortical hierarchy is comparatively shallow in the mouse (Harris et al. 2019), feedforward projections tend to arise more prominently from supragranular neurons, whereas feedback projections contain a larger infragranular contribution (Coogan and Burkhalter 1990, 1993; Berezovskii et al. 2011; Vezoli et al. 2021). VGluT1 mRNA is abundantly expressed in both supra- and infragranular layers of the adult mouse visual and somatosensory cortices, consistent with the predominance of VGluT1+ boutons in both feedforward and feedback projections (Nakamura *et al*. 2007). On the other hand, VGluT2 mRNA-expressing neurons have been reported primarily in layers 2/3, with smaller populations in infragranular cortical layers (Nakamura *et al*. 2007). Supragranular neurons are consequently plausible sources of the VGluT2+ boutons detected in feedforward corticocortical projections.

Although sparse, layer 4 corticocortical projection neurons do exist and may provide an additional source of these boutons. Such neurons have been identified in mouse cortex and exhibit laminar projection patterns characteristic of feedforward connections (Tamamaki and Tomioka 2010; Minamisawa et al. 2018; Tasic et al. 2018; Harris *et al*. 2019). However, neither the supragranular nor the layer 4 origin hypothesis can be resolved from the laminar position of the boutons alone. Establishing the cellular origin of VGluT2+ corticocortical boutons will require approaches combining layer- or cell-type-specific tracing with detection of the transporter phenotype.

### Corticofugal projections are predominantly associated with VGluT1

Corticofugal projections to the thalamus, superior colliculus, striatum, and pontine nuclei were strongly dominated by VGluT1. Among singly labelled VGluT+ boutons, corticothalamic and corticopontine boutons expressed VGluT1 exclusively, whereas VGluT1 accounted for more than 95% of the observed corticotectal and corticostriatal boutons. These findings are broadly consistent with comparative studies associating VGluT1 with descending projections of the visual cortex (Balaram, Hackett*, et al.* 2011; Balaram, Takahata*, et al.* 2011; Balaram *et al*. 2015; Abbas Farishta *et al*. 2022). Nevertheless, the rare VGluT2+ boutons detected in the corticostriatal and corticotectal pathways indicate that the molecular phenotype of cortical output projections is not invariably homogeneous.

#### Corticothalamic projections

All singly labelled boutons examined in projections from V1 and V2_L_ to the dLGN and LP were VGluT1+. Corticogeniculate feedback arises predominantly from layer 6, whereas projections to the higher-order LP nucleus originate from neurons in both layers 5 and 6 (Sherman 2017; Harris *et al*. 2019; de Souza et al. 2021). VGluT1 mRNA is widely expressed among neurons in both these infragranular layers, while VGluT2 mRNA is restricted to smaller neuronal populations and has been reported predominantly in layer 5 rather than layer 6 of the adult mice visual cortex (Nakamura *et al*. 2007). The uniform VGluT1 phenotype observed at identified corticogeniculate boutons is consistent with their predominant layer 6 origin and with comparative evidence associating corticogeniculate neurons with VGluT1 expression (Balaram *et al*. 2013; Balaram *et al*. 2015). More direct evidence was provided by Lindström *et al*. (2020), who demonstrated preferential VGluT1 expression and little VGluT2 colocalization at genetically identified layer 6 corticothalamic terminals in the mouse dLGN.

The transporter phenotype of corticopulvinar projections appears to vary across species, cortical areas, and laminar populations. In primates, the distributions of VGluT1 and VGluT2 mRNAs have led to the proposal that layer 6 corticopulvinar neurons predominantly express VGluT1, whereas some layer 5 neurons may contribute VGluT2-expressing projections (Balaram *et al*. 2013). In the tree shrew, VGluT1 is widely expressed in layers 5 and 6 of V1, while VGluT2 expression is restricted to smaller populations, suggesting that layer 6 corticopulvinar projections predominantly use VGluT1 and that some layer 5 projections may exhibit greater molecular heterogeneity (Balaram *et al*. 2015). Because these interpretations are based largely on the laminar distribution of transporter-expressing neurons rather than direct identification of corticopulvinar terminals, the correspondence between laminar origin and terminal phenotype remains inferential.

A similar interpretative limitation applies to the mouse. Projections from primary and extrastriate visual cortex to LP originate from neurons in deep layer 5 and layer 6, although the relative contributions of these populations vary among cortical areas and pathways (de Souza *et al*. 2021). Sparse VGluT2 mRNA-expressing neurons have also been reported in the deeper portion of layer 5 of the adult mice visual cortex (Nakamura *et al*. 2007). Whether these neurons belong to the population projecting to LP is unknown, but the overlap in their laminar distribution leaves open the possibility of a scarce VGluT2+ cortico-LP component. In the present study, however, no VGluT2+ boutons were detected in the V1- or V2_L_-to-LP projections. This finding may reflect a genuine predominance of VGluT1 in mouse cortico-LP pathways, the relatively sparse contribution of layer 5 neurons to the cortical projections labeled by the PHA-L, or limited sampling of a rare VGluT2+ component. The present results therefore demonstrate a uniform VGluT1 phenotype among the sampled cortico-LP boutons but may not establish that VGluT2 is entirely absent from these projections.

Moderate VGluT2 immunoreactivity was nevertheless present within LP, while substantially stronger labelling was observed in the adjacent dLGN. The strong VGluT2 signal in the dLGN is consistent with its dense retinal input, which forms VGluT2+ terminals (Fujiyama *et al*. 2003; Islam and Atoji 2009). By contrast, the moderate VGluT2 immunoreactivity observed in LP might likely originates mostly from non-cortical afferents, including the VGluT2+ tectothalamic projection identified in this study, however, a rare cortical contribution cannot be excluded.

#### Corticostriatal projections

Most boutons in the projections from V1- and V2_L_-to-striatum were VGluT1+, consistent with evidence that VGluT1+ and VGluT2+ terminals in the striatum originate respectively predominantly from cortical and thalamic inputs (Fujiyama et al. 2004). Indeed, glutamatergic afferents to the striatum originate from the cerebral cortex and the intralaminar thalamic nuclei (Smith and Bolam 1990), and both transporter isoforms are present in striatal terminals throughout development (Nakamura et al. 2005). The present results extend these regional observations by directly demonstrating that anatomically identified corticostriatal boutons arising from visual cortex are predominantly associated with VGluT1.

A small number of corticostriatal boutons expressed VGluT2, and one bouton in the V1 projection to striatum exhibited detectable VGluT1/VGluT2 colocalization. One possible explanation is that these rare boutons originate from cortical populations outside the layer 5 neurons that provide the principal corticostriatal output. In support of this possibility, sparse layer 4 pyramidal neurons have been shown to project to the dorsal striatum in mouse auditory cortex (Bertero et al. 2022), while VGluT2 expression and VGluT1/VGluT2 coexpression have been reported in subsets of neurons within layers 2–4 (De Gois et al. 2006; Nakamura *et al*. 2007). Layer 4 neurons could therefore represent a potential source of the rare VGluT2+ or double-labelled corticostriatal boutons detected here. Whether these terminals originating from cortical projection originating from layer 4 and or 5 remains to be investigated.

#### Corticotectal projections

Corticotectal projections from both V1 and V2_L_ were strongly dominated by VGluT1, with only a few VGluT2+ boutons detected in the superficial and intermediate layers of the superior colliculus. This pattern is consistent with regional and projection-based evidence reported across several mammalian lineages, including primates (Balaram, Hackett*, et al.* 2011; Balaram *et al*. 2013), tree shrews (Balaram *et al*. 2015), carnivores (Abbas Farishta *et al*. 2022) and rodents (Kaneko and Fujiyama 2002; Kaneko et al. 2002). Together, these comparative findings suggest that the predominance of VGluT1 in visual cortical projections to the superior colliculus is broadly shared across mammalian species. In contrast, the strong VGluT2 immunoreactivity of the superficial collicular layers has been attributed primarily to retinotectal and other subcortical afferents rather than to cortical input. By directly identifying boutons arising from V1 and V2_L_, the present study confirms that visual cortical afferents are predominantly VGluT1+ and therefore contribute little to the strong regional VGluT2 signal observed in the superficial superior colliculus.

Although this general organization accounts for the predominant VGluT1 phenotype of corticotectal boutons, it does not explain the small VGluT2-positive component identified here. Corticotectal projections originate mainly from layer 5 neurons, where VGluT1 mRNA is widely expressed but VGluT2 mRNA also occurs in smaller neuronal populations in the mouse visual cortex (Nakamura *et al*. 2007). These less numerous VGluT2-expressing layer 5 neurons constitute a possible source of the rare VGluT2+ corticotectal boutons observed here. However, because transporter expression was assessed at terminals rather than in their associated cell bodies, the laminar and cellular origins of these boutons remain to be established.

#### Corticopontine projections

All singly labelled corticopontine boutons arising from V1 and V2_L_ were VGluT1-positive. Both VGluT1 and VGluT2 immunoreactivities have previously been described in the pontine nuclei (Bellocchio et al. 1998; Kaneko and Fujiyama 2002; Kaneko *et al*. 2002; Varoqui *et al*. 2002), with VGluT1 being particularly prominent relative to its expression in many other brainstem nuclei (Kaneko *et al*. 2002). Based on this regional distribution and the predominance of VGluT1 in cortical projection systems, corticopontine afferents have been proposed as a major source of VGluT1-positive terminals in the pons (Kaneko and Fujiyama 2002; Kaneko *et al*. 2002). The marked reduction in high-affinity glutamate uptake following transection of the cerebral peduncles provides additional evidence for a substantial glutamatergic cortical input to the pontine nuclei (Thangnipon et al. 1983). By directly identifying boutons arising from V1 and V2_L_, the present results confirm that sampled visual corticopontine projections contribute a predominantly VGluT1-positive input to the pons.

### Thalamic and tectal projections are predominantly associated with VGluT2

Thalamic and tectal projections were strongly dominated by VGluT2. Nearly all singly labelled thalamocortical and tectothalamic boutons were VGluT2+, as were all sampled LP-to-striatum and tectopontine boutons. Additional VGluT1/VGluT2 double-labelled boutons were detected in the dLGN-to-V1, SC-to-dLGN, and SC-to-LP projections. Thus, VGluT2 represented the predominant phenotype of both thalamic and tectal outputs, although rare dual expression indicated that the two transporter phenotypes were not invariably mutually exclusive.

#### Thalamocortical projections

VGluT2-positive boutons predominated in thalamocortical projections arising from both the dLGN and LP. This finding is consistent with comparative studies associating ascending thalamocortical projections with VGluT2 across mammalian visual systems (Balaram, Takahata*, et al.* 2011; Balaram *et al*. 2013; Balaram *et al*. 2015; Abbas Farishta *et al*. 2022). More direct evidence has demonstrated VGluT2 expression at anatomically identified geniculocortical terminals in tree shrews (Familtsev et al. 2016), mice (Coleman et al. 2010), ferrets (Nahmani and Erisir 2005), and primates, as well as in pulvinar projections to the primate cortex (Marion *et al*. 2013).

In the dLGN-to-V1 projection, almost all singly labelled boutons were VGluT2-positive and were particularly abundant in layer 4, although they were also detected in supra- and infragranular layers. Four additional boutons in layer 4 exhibited detectable VGluT1/VGluT2 colocalization. Previous studies across rodents, primates, and tree shrews have consistently reported strong VGluT2 expression together with lower levels of VGluT1 in the dLGN (Fremeau *et al*. 2001; Fujiyama et al. 2001; Herzog *et al*. 2001; Balaram, Hackett*, et al.* 2011; Balaram, Takahata*, et al.* 2011; Balaram *et al*. 2013; Balaram *et al*. 2015). Coexpression of the two transcripts has been demonstrated directly in subsets of thalamic relay neurons, including neurons of the adult mouse dLGN (Herzog *et al*. 2001; De Gois et al. 2005; Barroso-Chinea et al. 2007; Nakamura *et al*. 2007). The present findings extend this evidence to anatomically identified terminals by demonstrating that a small number of mice geniculocortical boutons contain both transporter isoforms.

Developmental evidence further contextualizes the rarity of this phenotype. VGluT1/VGluT2 colocalization at neocortical terminals is transiently elevated during early postnatal development and remains more frequent in somatosensory than in visual cortex in adulthood (Nakamura *et al*. 2005; Nakamura *et al*. 2007). The small number of double-labelled dLGN boutons identified in adult V1 is consistent with this regional and developmental pattern, although the present analysis cannot determine whether the two transporters occupy the same synaptic vesicles within individual boutons.

All singly labelled boutons examined in cortical projections from LP were VGluT2+. These boutons were distributed across multiple cortical layers, with substantial representation in layers 1 and 4. Strong VGluT2 mRNA expression has been reported in the rodent LP and primate pulvinar, consistent with the predominance of VGluT2 in their cortical projections (Barroso-Chinea *et al*. 2007; Nakamura *et al*. 2007; Balaram, Takahata*, et al.* 2011; Balaram *et al*. 2013). In the tree shrew pulvinar, both transcripts are present, although VGluT2 expression generally exceeds that of VGluT1, leading to the proposal that some pulvinocortical projections may use both isoforms (Balaram *et al*. 2015). No VGluT1-positive or double-labelled LP-to-cortex boutons were detected in the present sample. These findings indicate that VGluT2 might be the predominant transporter in mouse LP thalamocortical projections

#### Thalamostriatal projections

All six boutons sampled from the LP-to-striatum projection were VGluT2+. Although the major thalamostriatal inputs arise from intralaminar and motor-related thalamic nuclei, an LP-to-striatum projection has been demonstrated in the mouse (Berendse and Groenewegen 1990; Elena Erro et al. 2002; Pan et al. 2010). Pulvinar projections to the striatum have also been reported across several other mammalian species, including tree shrews (Lin et al. 1984; Day-Brown et al. 2010), carnivores (Beckstead 1984; Takada, Itoh, Sugimoto, et al. 1985; Takada, Itoh, Yasui, et al. 1985; Harting et al. 2001), squirrels (Lin *et al*. 1984) and tenrec (Kunzle 2006).

The VGluT2 phenotype of the sampled LP-to-striatum boutons is consistent with previous evidence associating thalamostriatal terminals with VGluT2 (Bacci et al. 2004; Hur and Zaborszky 2005; Aymerich et al. 2006). VGluT2 is the predominant transporter expressed by midline and intralaminar thalamostriatal neurons, whereas some sensory thalamic nuclei contain neuronal populations expressing both VGluT1 and VGluT2 transcripts (Barroso-Chinea *et al*. 2008). In LP, VGluT2 mRNA expression is moderate to strong, while VGluT1 mRNA expression is comparatively weak (Barroso-Chinea *et al*. 2007). These observations are compatible with the phenotype identified in the present study.

#### Tectothalamic projections

VGluT2+ boutons accounted for nearly all singly labelled boutons in the SC-to-dLGN and SC-to-LP projections. Six additional tectothalamic boutons exhibited VGluT1/VGluT2 colocalization, including three in each projection. The superior colliculus provides a major component of the extrageniculate visual pathway to LP or its pulvinar homologues across mammals and also projects to the dLGN in the mouse and several other species. Although the SC-to-dLGN and SC-to-LP pathways arise from at least partly distinct neuronal populations (Gale and Murphy 2018), their transporter phenotypes were similarly predominant by VGluT2.

This predominance is consistent with the strong expression of VGluT2 and comparatively weak expression of VGluT1 reported in the superior colliculus of rodents (Kaneko *et al*. 2002) and primates (Balaram, Hackett*, et al.* 2011; Balaram, Takahata*, et al.* 2011). Similar distributions in tree shrews and carnivores have led to the proposal that tectopulvinar projections predominantly use VGluT2 (Balaram *et al*. 2015; Abbas Farishta *et al*. 2022). In primates, VGluT2 mRNA-expressing neurons are particularly prominent in the lower superficial grey layers that contribute to pulvinar projections, whereas VGluT1 expression is absent or weak in these layers (Balaram *et al*. 2013). The present findings provide direct terminal-level evidence that VGluT2 is likewise the principal transporter in mouse tectothalamic projections.

The detection of a small number of double-labelled boutons nevertheless qualifies proposals of an exclusive VGluT2+ tectothalamic phenotype. These boutons could arise from restricted collicular populations that express low levels of VGluT1 in addition to VGluT2. However, because the present analysis did not identify the parent neurons of individual boutons, it remains unknown whether the double-labelled SC-to-dLGN and SC-to-LP terminals originate from the same or distinct collicular cell types. Their functional significance also remains uncertain given their low prevalence.

#### Tectopontine projections

The predominance of VGluT2 in tectal efferents extended beyond the ascending tectothalamic pathways. All singly labelled tectopontine boutons sampled following superior colliculus injections were VGluT2+. This phenotype contrasted directly with that of visual corticopontine projections, in which all singly labelled boutons arising from V1 and V2_L_ were VGluT1+. Thus, the visual cortex and superior colliculus provide molecularly distinct glutamatergic inputs to the pontine nuclei despite both pathways being anatomically descending. This complementary organization demonstrates that VGluT phenotype is more closely associated with neuronal origin and pathway identity than with projection direction alone. It may also explain, at least in part, the coexistence of VGluT1 and VGluT2 immunoreactivities previously reported within the pontine nuclei (Bellocchio *et al*. 1998; Kaneko and Fujiyama 2002; Kaneko *et al*. 2002; Varoqui *et al*. 2002).

Visual corticopontine and tectopontine pathways constitute parallel routes through which cortical and collicular signals can access pontocerebellar circuits, as demonstrated anatomically in primates (Glickstein et al. 1990). Within this dual-input organization, the rodent corticopontine pathway is spatially structured, cortical afferents form topographically organized terminal fields, with occipital inputs preferentially targeting lateral and rostral pontine regions (Leergaard and Bjaalie 2007). The present findings add a molecular dimension to this anatomical organization by showing that visual corticopontine boutons predominantly express VGluT1, whereas tectopontine boutons express VGluT2. The two pathways therefore constitute anatomically distinct and molecularly differentiated channels for conveying visual information to the pons.

The tectopontine pathway is unlikely to be the only source of VGluT2+ terminals in the pontine nuclei, which receive numerous glutamatergic afferents from other subcortical structures (Aas 1989; Mihailoff et al. 1989; Border and Mihailoff 1991; Giolli et al. 2001). Nevertheless, the present study directly identifies the superior colliculus as one such source. Together with the

VGluT1 phenotype of identified corticopontine boutons, this finding provides a projection-specific anatomical basis for the presence of both transporter isoforms within the pons.

### Terminal morphology depends on VGluT phenotype and projection direction

Bouton morphology in the adult mouse visual system was not explained by VGluT phenotype alone. Bouton area and VGluT-immunoreactive punctum area each showed an interaction between transporter isoform and projection direction. Within ascending projections, VGluT2+ boutons and puncta were larger than their VGluT1+ counterparts, whereas within descending projections the bouton-area difference was nonsignificant and VGluT1+ puncta were modestly larger. Moreover, VGluT1+ boutons were smaller in ascending than in descending pathways, while VGluT2+ bouton area remained comparatively stable across directions. Terminal morphology therefore appears to depend on the combination of transporter phenotype and pathway identity rather than on VGluT isoform alone.

The larger VGluT2+ boutons observed within ascending pathways are consistent with previous findings in the visual system. In ferret V1, VGluT2+ layer 4 terminals corresponded morphometrically to anatomically identified thalamocortical terminals and were larger than unlabelled terminals, formed longer synaptic contacts, and more frequently innervated dendritic spines (Nahmani and Erisir 2005). In rat retinorecipient structures, immunoelectron microscopy similarly showed that VGluT2+ retinal terminals were larger than VGluT1+ terminals in the dLGN and superior colliculus (Fujiyama *et al*. 2003). These studies support an association between VGluT2 expression and large terminal profiles in selected ascending sensory pathways, but they do not establish that VGluT2 intrinsically determines bouton size.

Evidence from other circuits further argues against a fixed relationship between VGluT isoform and terminal size. In the macaque thalamus, giant presumed driver terminals can express either VGluT1 or VGluT2 depending on their cortical or subcortical origin (Rovo *et al*. 2012). The expected size relationship is even reversed in the rat striatum, where VGluT1+ corticostriatal axospinous terminals are larger than VGluT2+ thalamostriatal terminals and more frequently form synapses associated with perforated postsynaptic densities (Liu et al. 2011). Substantial variation also occurs among cortical outputs themselves. In the mouse posterior thalamic nucleus, layer 6b corticothalamic varicosities are markedly smaller and structurally simpler than most layer 5 varicosities (Hoerder-Suabedissen et al. 2018). Similarly, anatomically identified cortical projections can form either giant, driver-like terminals, as in the piriform-to-mediodorsal thalamic pathway (Pelzer et al. 2017), or predominantly small terminals, as in the auditory corticocollicular pathway (Nakamoto et al. 2013). Together with the present results, these observations indicate that neuronal origin, laminar source, target structure, and local synaptic organization influence terminal size more strongly than transporter phenotype alone.

The punctum-area findings should be interpreted more cautiously. VGluT2 immunoreactivity is concentrated within vesicle-containing thalamocortical boutons and is largely absent from intervening axonal segments, supporting its use to delineate presynaptic terminal compartments (Nahmani and Erisir 2005; Coleman *et al*. 2010). Large anatomically identified thalamocortical boutons may contain multiple synapses, extensive vesicle pools, large postsynaptic densities, and several mitochondria, features that plausibly contribute to their transmission efficacy (Rodriguez-Moreno et al. 2018). The punctum measurements therefore provide complementary evidence that VGluT-defined terminal populations differ across pathway classes, but not a direct functional classification. Overall, the convergence of VGluT2 expression and larger boutons within ascending projections is compatible with a driver-like organization, whereas the pathway-specific exceptions argue against a one-to-one correspondence among VGluT isoform, terminal size, and driver–modulator function.

## Conclusion

VGluT1 and VGluT2 exhibited a strong but non-absolute projection-specific organization across the mouse visual system. VGluT1 predominated in cortical feedback and corticofugal pathways, whereas VGluT2 characterized most thalamic and tectal outputs. However, the predominance of VGluT1 in ascending corticocortical projections and of VGluT2 in the descending tectopontine pathway demonstrates that transporter phenotype is more closely associated with neuronal origin and pathway identity than with projection direction alone. Terminal morphology was similarly context dependent: VGluT2+ boutons and puncta were larger than their VGluT1+ counterparts within ascending projections, but this relationship was not generalized across descending pathways. The rare detection of VGluT1/VGluT2 colocalization further indicates that the two transporter phenotypes are not invariably mutually exclusive. VGluT isoforms therefore provide informative markers of visual pathway organization, but neither isoform identity nor terminal size alone establishes projection direction or driver–modulator function. Pathway-specific electrophysiological and ultrastructural analyses will be required to determine the functional significance of these molecular and morphological phenotypes.

## Acknowledgements

The authors thank the animal care staff of the Université du Québec à Trois-Rivières for their assistance with animal husbandry and welfare.

## Author Contributions

GL: Formal analysis; Visualization; Writing – original draft; Writing – review and editing. RTL: Investigation; Data curation; Formal analysis; Visualization. DB: Conceptualization; Funding acquisition; Supervision; Writing – original draft; Writing – review and editing. All authors reviewed and approved the final version of the manuscript and agree to be accountable for the work.

## Funding Information

This work was supported by grants from the Natural Sciences and Engineering Research Council of Canada awarded to Denis Boire for the projects “Cortical feedback microcircuits in the visual system of the mouse” (Grant No. 203702-2013) and “Linking cortical circuits to subcortical output structure” (Grant No. 522474-2018).

## Conflict of Interest Statement

The authors declare that they have no conflicts of interest.

## Ethics Statement

All experimental procedures were approved by the Comité de bons soins aux animaux of the Université du Québec à Trois-Rivières (protocol no. DB8) and were conducted in accordance with the guidelines of the Canadian Council on Animal Care.

## Data Availability Statement

The data supporting the findings of this study and the R code used for the statistical analyses are available from the corresponding author upon reasonable request.

## Notes

### Competing Interest Statement

The authors have declared no competing interest.

